# Targeted inhibition of BET proteins in HPV-16 associated head and neck squamous cell carcinoma reveals heterogeneous transcription response

**DOI:** 10.1101/2023.10.02.560587

**Authors:** Aakarsha Rao, Zijian Ni, Dhruthi Suresh, Chitrasen Mohanty, Albert R. Wang, Denis L Lee, Kwangok P. Nickel, Sooryanarayana Varambally, Randall J. Kimple, Paul F. Lambert, Christina Kendziorski, Gopal Iyer

## Abstract

Integrated human papillomavirus (HPV-16) associated head and neck squamous cell carcinoma (HNSCC) tumors have worse survival outcomes compared to episomal HPV-16 HNSCC tumors. Therefore, there is a need to differentiate treatment for HPV-16 integrated HNSCC from other viral forms. We analyzed TCGA data and found that HPV+ HNSCC expressed higher transcript levels of the bromodomain and extra terminal domain (BET) family of transcriptional coregulators. However, the mechanism of BET protein-mediated transcription of viral-cellular genes in the integrated viral-HNSCC genomes needs to be better understood. We show that BET inhibition downregulates E6 significantly independent of the viral transcription factor, E2, and there was overall heterogeneity in the downregulation of viral transcription in response to the effects of BET inhibition across HPV-associated cell lines. Chemical BET inhibition was phenocopied with the knockdown of BRD4 and mirrored downregulation of viral E6 and E7 expression. Strikingly, there was heterogeneity in the reactivation of p53 levels despite E6 downregulation, while E7 downregulation did not alter Rb levels significantly. We identified that BET inhibition directly downregulated c-Myc and E2F expression and induced CDKN1A expression. Overall, our studies show that BET inhibition provokes a G1-cell cycle arrest with apoptotic activity and suggests that BET inhibition regulates both viral and cellular gene expression in HPV-associated HNSCC.

## Introduction

Papillomaviruses, a diverse group of non-enveloped, double-stranded DNA viruses, are known for their specific affinity for squamous epithelial tissues across various host species (1, 2). Their life cycle is intricately intertwined with the differentiation program of host epithelial cells, commencing with an initial infection of basal epithelial cells and culminating in a productive phase as these cells differentiate and migrate toward the surface, ultimately releasing new virions without causing cell lysis (3–7).

The medical significance of papillomaviruses is substantial, as they are implicated in a spectrum of benign conditions such as warts. However, their more critical association with malignancies, including cervical, anal, and oropharyngeal cancers, underscores the profound impact of these viruses on human health (4, 8, 9). Despite the introduction of vaccines targeting high-risk papillomavirus types, which has significantly reduced the incidence of associated diseases, global disparities in vaccine accessibility persist. Moreover, a comprehensive understanding of the initial phases of the viral life cycle, particularly the establishment of infection and persistence, holds the potential for innovative therapeutic interventions capable of impeding progression toward malignancy.

Human papillomaviruses (HPV) possess a compact genome commonly encoding eight proteins, categorized as early (E) and late (L) proteins. Among the early proteins, E1 and E2 play pivotal roles in viral DNA replication, with E1 serving as a DNA helicase (10) and E2 regulating transcription while acting as a segregation factor during cell division (11). E4, often considered a link between early and late gene expression, assumes roles in virus release and can disrupt the host cell’s intermediate filament network (12). The smallest HPV protein, E5, enhances the proliferation of infected cells and contributes to immune evasion by downregulating MHC class I molecules (13, 14) . The notorious oncogenes, E6 and E7, hold profound implications for healthcare: E6 targets p53 for degradation (15, 16), while E7 binds to the retinoblastoma protein (Rb), leading to cell cycle dysregulation (17, 18) .

The HPV genome can exist in both episomal (extrachromosomal) and integrated forms. While the episomal form is essential for the productive viral life cycle, integration into the host genome is often associated with cellular transformation, making this process pivotal in HPV-induced carcinogenesis. The late proteins, L1 and L2, constitute the viral capsid, with L1 being employed in vaccine formulations(19) . A comprehensive understanding of these viral proteins and the virus’s genomic state provides invaluable insights into HPV’s pathogenesis and its impact on host cells.

While the majority of HPV infections are benign and self-limiting, persistent infections by high-risk HPV types are etiologically linked to various cancers. Cervix, oropharyngeal, anal, penile, vulval, and vaginal tissues can undergo malignant progression. In the initial stages of infection, HPV exists as an episome—a circular, extrachromosomal DNA(20). This form enables the virus to maintain low-level replication without causing cellular damage, often leading to transient infections. During carcinogenesis, HPV DNA can integrate into the host genome (21). Integration often disrupts the E2 gene, leading to increased expression of E6 and E7 oncogenes, propelling the cell toward malignancy. However, not all integrated HPV genomes are disrupted, as recent studies have shown that some cancers harbor non-disrupted, integrated HPV genomes, suggesting diverse mechanisms of HPV-mediated oncogenesis (22).

Although E6 and E7 are crucial in HPV-induced carcinogenesis, they are insufficient to cause cancer; additional genomic alterations and co-factors are typically required. Therapeutic interventions targeting E6 and E7 proteins, foreign to human cells and consistently expressed in HPV-positive tumors, represent ideal therapeutic targets. Agents inhibiting these proteins or inducing their degradation are under investigation. Therapeutic vaccines aim to enhance the immune response against E6 and E7-expressing cells, attempting to eliminate HPV-transformed cells before malignancy ensues. Given E6’s impact on p53 and E7’s on Rb, therapies restoring these pathways, such as MDM2 inhibitors, hold promise against HPV-associated cancers. Considering that HPV integration can lead to epigenetic alterations, drugs modifying the epigenetic landscape, such as DNA methyltransferase inhibitors or histone deacetylase inhibitors, may offer therapeutic potential. A recent and promising area of therapeutic intervention is the class of bromodomain and extra terminal (BET) inhibitors, which target proteins containing bromodomains, including BRD4. BRD4, a member of the BET family, plays pivotal roles in normal physiological processes, particularly in regulating gene transcription. By recognizing acetylated histones (22), BRD4 facilitates the recruitment of positive transcription elongation factor b (P-TEFb) to chromatin (23) thus promoting the elongation phase of transcription and influencing cell cycle progression (24–26), DNA damage response (27–31), and cellular differentiation (32).

BRD4’s interaction with episomal HPV genomes is well-documented. It interacts with the viral E2 protein, essential for maintaining the episomal state of the HPV genome (34). This interaction is crucial for episomal HPV genome segregation during cell division and the regulation of viral transcription (35). However, once HPV integrates into the host genome, the E2 gene is often disrupted, leading to deregulated expression of E6 and E7 oncogenes, which are critical for the oncogenic potential of high-risk HPVs. Although the canonical role of BRD4 in interacting with E2 is altered upon viral genome integration, BRD4 still plays a role in the transcriptional regulation of HPV genes in the integrated state due to its broader roles in chromatin organization and gene transcription through its association with enhancers and super-enhancers. For instance, McBride et al. demonstrated that in cervical neoplastic cell lines, such as W12 clone 20861, tandemly integrated HPV gives rise to super enhancers. In these cases, inhibiting BRD4 has been shown to impact the expression of key viral genes E6 and E7 (33, 34). However, it is crucial to acknowledge that not all viral integrations will culminate in super enhancer formation. The specific outcome hinges on various factors, including the nature of the virus, the integration site within the host genome, and the characteristics of the host cell. Furthermore, super enhancer formation tends to be closely linked to specific cellular programs and cell types (35). Consequently, the context in which viral integration occurs remains pivotal.

In the context of HPV-associated head and neck squamous cell carcinoma (HNSCC), the impact of BRD4 on super enhancers that have dispersed viral genomes are not yet fully elucidated. While BRD4’s conventional role in maintaining episomal HPV genomes through E2 interaction may be compromised upon viral integration, its broader involvement in cellular transcription suggests it may still exert influence over the transcriptional activity of integrated HPV genomes. This realization underscores the potential therapeutic value of targeting BRD4 and BET proteins, not only for episomal HPV but also for integrated forms of the virus. This could prove particularly beneficial in the treatment of cervical and head and neck cancers, which originate from distinct anatomical sites.

While BRD4 regulation in cervical cancer lines has been described extensively(33, 36), very little is known in the context of integrated HPV HNSCC tumors. HPV HNSCC patients with HPV16-associated head and neck squamous cell carcinoma (HNSCC), the presence of integrated HPV-16 genomic forms correlates with more unfavorable survival outcomes (37) compared to their episomal counterparts (38). This study explored the effects of the pan-BET inhibitor JQ1 on viral transcription in seven head and neck squamous cell carcinoma (HNSCC) cell lines associated with HPV16. These cell lines possess integrated HPV16 DNA in their genomes, and this integration process reflects the complexity of how HPV contributes to cancer development. While the HPV proteins E6 and E7 are pivotal in driving carcinogenesis, additional genetic changes are typically required for cancer progression. Therefore, the cell lines we used not only contained integrated HPV DNA but also exhibit various genetic alterations. This unique combination allowed us to investigate the intricate relationship between BET protein activity and the regulation of both viral and cellular gene expression when we inhibited BET proteins with JQ1. JQ1 treatment emerged as a potent regulator, significantly inhibiting the expression of the E6 oncogene to a greater extent than E7 in these cell lines. Furthermore, we unveiled an additional facet of JQ1’s action—its role in repressing E2 expression in a subset of these cell lines, which was, surprisingly, expressed significantly. Another finding was JQ1-induced downregulation of c-MYC across all the cell lines.

This finding was particularly intriguing because, unlike in lung adenoma-carcinoma cell lines (39), the sensitivity to JQ1 in our HPV-associated HNSCC cell lines was not dependent on c-MYC expression. Moreover, our study revealed a crucial consequence of E6 downregulation, leading to the reactivation of the tumor suppressor p53. In stark contrast, downregulation of E7 did not alter the levels of the retinoblastoma protein (RB). The combined effects of JQ1 treatment on E6 and E7 expression culminated in a substantial increase in p21 expression, further underscoring the potential interplay between E6-p53 and E7-Rb with p21 expression. Lastly, we unraveled a complex circuitry of JQ1 treatments, revealing that JQ1 disrupted the classical Rb-E2F complex governing the G1-S transition, primarily through the transcriptional repression of E2F. This intricate network of interactions illuminated numerous promising therapeutic intervention points within the realm of integrated HPV-associated HNSCC.

## Materials and Methods

### Cell culture

UD: SCC2, UM: SCC47, UPCI: SCC90, UM: SCC104, 93VU147, UPCI: SCC152, UPCI: SCC154 were obtained from NCI-Head and Neck SPORE biobank (University of Wisconsin-Madison). Cell lines were cultured in Dulbecco’s Modified Eagle Medium 4.5g/L glucose, L-glutamine and sodium pyruvate (DMEM) supplemented with 10% fetal bovine serum (FBS) and maintained in 5% (vol/vol) CO_2_ and 95% (vol/vol) air in a humidified incubator at 37°C. All cell lines are tested negative for mycoplasma, and STR tested by LabCorp.

### Southern blot

Assessment of the integration status of the HPV-positive HNSCC cell lines was characterized for each cell line and digested with 7 µg of total genomic DNA at 37°C overnight (20 hours) with either *EcoRV* (New England BioLabs) or *Bam HI.* HPV-positive HNSCC samples and λ HindIII marker and standards were loaded on a 1X TAE gel apparatus for gel electrophoresis at 30V overnight, followed by EtBr stain and destaining. The gel underwent denaturation and neutralization washes and was transferred overnight (20 hours) to a Hybond membrane (Amersham). The membrane was subjected to UV crosslink with a Strata Linker on auto crosslink. The ^32^P radiolabeled HPV16 genotype-specific single-stranded oligonucleotides probe was incubated in a Techne hybridization tube in a Techne hybridizer HB-1D incubator at 48°C. Membrane was subsequently washed with Church hybridization buffer followed by exposure on a Molecular Dynamics cassette overnight (24 hours). The radio-labelled blot was imaged with a Typhoon 8610 imager (Molecular Dynamics).

### Cell Viability Assays

All head and neck cancer cell lines were treated with dimethyl sulfoxide (DMSO)(vehicle), 0.5 μM (+)-JQ1 (APExBIO) or (-)-JQ1 (APExBIO) for 96 hours. The numbers of live cells in each condition were counted using Bio-rad TC20 digital cell counter and trypan blue exclusion. The cell numbers were then normalized to the vehicle-treated group for all experiments.

### Pan-BET inhibitor treatments and cell viability determination

To determine the half maximal inhibitory concentrations (IC50s) for each head and neck cell cancer line, HNSCC cells were plated into a 96-well plate at 1500-4500 cells/well and treated with various (+)-JQ1 concentrations (range from 10 nM to 10 μM) for 96 h. To measure cell viability, 10 µL of Prestoblue Cell Viability Reagent (Thermo Fisher) was added to each well (100 µL of medium) and incubated at 37 °C for 1 and 4 h. The plates were read by CLARIOstar microplate reader (BMG Labtech, Software version: 5.21 R2, Firmware version: 1.15) with the excitation/emission at 535/590 nm. The raw absorbance values were first subtracted by the average absorbance value of the background control (medium only). Then, the cell survival rate at each (+)-JQ1 concentration was calculated by normalizing to the vehicle control signal (DMSO-treated cells). The data was curve fitted using the sigmoidal dose-response equation (Y= Bottom + (Top-Bottom)/(1+10^((LogIC_50_-X)*HillSlope)) in OriginLab software (OriginLab) to determine the IC_50_ values.

### Lentiviral transduction

The MISSION lentiviral-based shRNA vector collections from Sigma Aldrich (St. Louis, MO, USA) were used for long-term silencing of BRD4. Briefly, 1.6×10^4^ cells were cultured in 96-well plates, incubated for 18-20 h. Hexadimethrine bromide (8 μg/ml) was added after media removal, and effective multiplicity of infection (MOI) of lentiviral particles was established. [TRCN0000199427] [TRCN0000318771] and (V-591) [TRCN0000196576] were used to identify knockdown clones. After 18-20 h incubation at 37°C, puromycin was added at 1.0 µg/ml, and resistant clones were selected and grown.

### Analysis of Caspase 3/7 Activity – Apoptosis Assay

All cell lines were cultured in high-glucose DMEM complete with 10% FBS and in a 5% CO_2_ enriched humidified incubator at 37°C until 90% confluency. The cells were counted using Trypan Blue live/dead staining and were subsequently plated at a density of 3000 cells/well in a 96-well plate (TPP). They were allowed to adhere to the plate overnight, following which the cells were treated with 500nM (+)-JQ1 and (-)-JQ1. Post 24 hours of treatment, 100 µl/well of room temperature Caspase Glo 3/7 (Promega) reagent was added to the treated and control cells. After a 1hr incubation at room temperature, luminescence was measured using the CLARIOstar microplate reader (BMG Labtech, Software version: 5.21 R2, Firmware version: 1.15).

### Quantitative RT-PCR

All cell lines were pre-treated with 0.5 µM (+)-JQ1 or JQ1 (-)-(vehicle control) for 30 min and 24 h for RNA extraction. Samples were collected with Trizol reagent at 30 minutes and 24 hours post-treatment, extracted with chloroform, ethanol wash, phase-lock separation, and DNase I treatments were used to isolate RNA, and a high-capacity RNA-to-cDNA kit (Applied Biosystems) was used to synthesize first-strand cDNA. Quantitative PCR reactions were set up using PowerUp SYBR Green Master Mix (Applied Biosystems) and performed on a Bio-Rad CFX96 machine. Primers are listed in Table S1. Ct values were determined from the software CFX Maestro 1.1 (Bio-Rad), and the delta-delta Ct method was used to calculate the relative mRNA expressions of each target relative to the housekeeping gene(40).

**Table S1.**
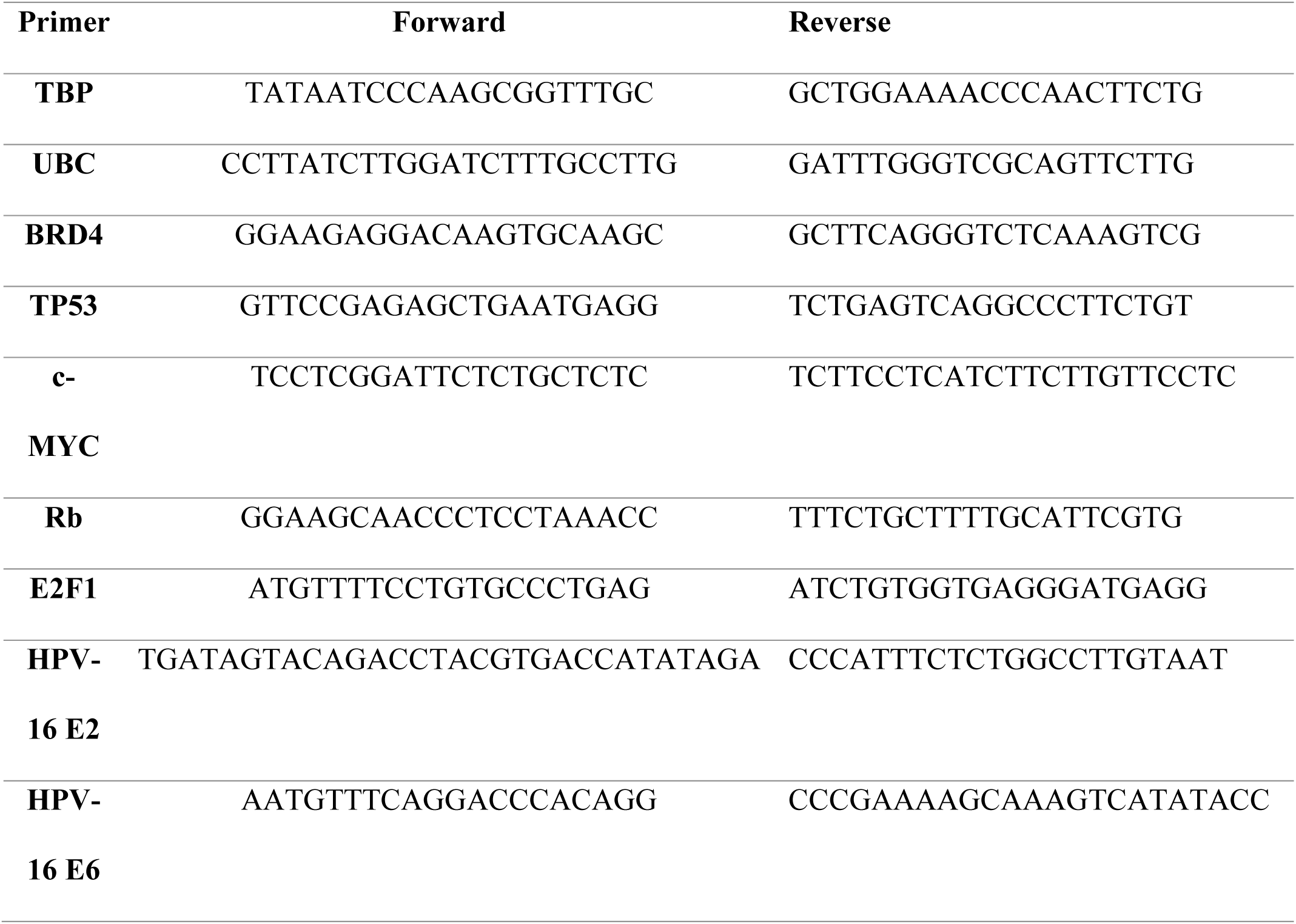
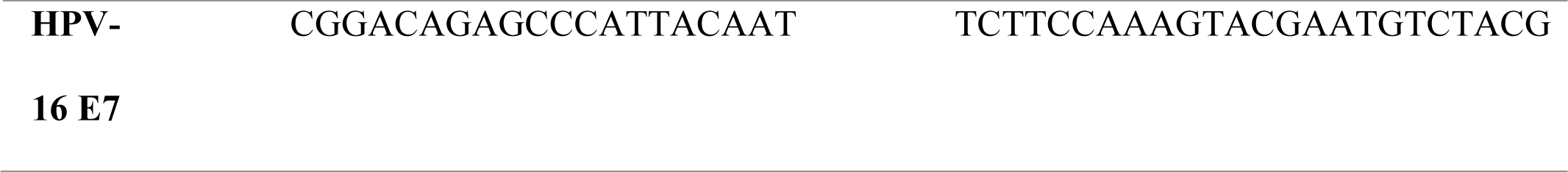

### Flow Cytometry

All cell lines were cultured in high-glucose DMEM complete with 10% FBS in a 5% CO_2_ enriched humidified incubator at 37°C. They were plated at a density of 0.5×10^6^ cells per 10cm dish. (+)-JQ1 was added to cells at 0.5µM concentration for 30 min, 24 and 48 hours to test its effect on cell cycle progression. DMSO was used as the vehicle control for the experiment since (+)-JQ1 was dissolved in the DMSO solvent. Cells were then harvested using Trypsin-EDTA (Corning) after incubation for 30 min, 24 and 48 hours. This was followed by rinsing cells with 1X PBS and fixing them in 70% EtOH (41). Following protocol (41), cells were stained with propidium iodide (PI) (0.5mg/ml, BioLegend) in a staining buffer containing PBS, DNA-free RNase (10mg/ml), 1% Triton-X (Sigma), and autoclaved water. Progression through the cell cycle was studied using Thermo Fisher Attune Flow Cytometer (4-laser, 14-color cytometer) with FSC=90V, SSC=280V, and PI=290V (yellow YL2 channel). The data was analyzed, and histograms were generated using ModFit (ModFit LT V5.0.9) software, and the bar graphs were prepared using Originlab software.

### Cell Lysate Preparation and Western Blot

<u>W</u>hole cell lysates were prepared by dissolving (10 million cells per treatment) cell pellets in RIPA buffer (G-Bioscience,) complete with 1X Protease and Phosphatase inhibitor cocktail (Thermo Scientific) and 1X Phosphatase inhibitor cocktails 1and 2 (Apex Bio). Cells were sonicated at 10% amplitude for 10 seconds using a sonicator (Fisher Scientific, Sonic dismembrator, Model 500). After a 30-minute incubation on a rotator at 4^0^C, cell samples were spun down at very high speed – 14,000 rpm for 15 mins at 4°C. Samples were diluted using 4X Laemlli sample buffer (Bio-Rad,) complete with 2-Mercaptoethanol and DI-water to bring all samples to a uniform concentration of 2µg/ul after protein quantitation using BCA assay. NuPage 4-12% Bis-Tris 26-well Midi gels (Invitrogen) were used to run the samples. The running buffer used was 1X MES Buffer made from 20X MES buffer. 30ug of protein was loaded on to each well for all the runs. The ladders used were Novex Sharp Pre-Stained Protein Standard and SeeBlue Plus2 Pre-Stained Protein Standard (Life Technologies). The gels were transferred on to a 0.2µm nitrocellulose membrane (Thermo-Scientific) by semi-dry transfer technique using the Bio-Rad Trans-Blot Turbo Transfer System protocols. All membranes were stained with Ponceau stain made fresh in the lab to ensure protein transfer. All membranes were blocked with 3% Bovine Serum Albumin (BSA) – Fraction V, Heat-shock treated (Fisher Scientific) with 0.5% sodium azide to avoid any possible contamination to the blot. Super signal west femto maximum sensitivity Substrate (Thermo Scientific) was used as the substrate. The blots were imaged using the Li-Cor F_C_ machine. Images were analysed and quantified using the Image Studio software (version 4.0.21). Each sample was normalized to its respective housekeeping loading control from the same day. This normalized value for each treatment was further normalized to the DMSO control to attain a “fold over control” expression of the protein under the stimulus of the drug. Western blot analysis followed standard protocol with indicated antibodies listed in Table below.

The primary antibodies used to probe the blots are as mentioned below:

| <u>Antibody</u> | <u>Host</u> | <u>Company</u> | <u>Reference #</u> |
| --- | --- | --- | --- |
| B-Actin | Mouse | Cell Signaling<br>Technology | CST 3700S |
| BRD4 | Rabbit | Abcam | Ab128874 |
| CDKN2A (p21) | Rabbit | Cell Signaling<br>Technology | CST 2947 |
| E2F1 | Mouse | Santa-Cruz | sc-251 |
| GAPDH | HRP-Linked | Proteintech | HRP60004 |
| Myc | Rabbit | Santa-Cruz | sc-40 |
| Total P53 | Rabbit | Cell Signaling<br>Technology | CST 2527 |
| Total Rb | Rabbit | Cell Signaling<br>Technology | CST 9313 |
| P-Rb (S780) | Rabbit | Sigma Aldrich | R6275 |
| P-Rb (S795) | Rabbit | Cell Signaling<br>Technology | CST 9301 |
| P-Rb (S807/S811) | Rabbit | Cell Signaling<br>Technology | CST 9308 |

**Secondary Antibody and image development:**
| Antibody | Dilution used | Company | Reference # |
| --- | --- | --- | --- |
| Mouse IgG | 1:5000 | Cell Signaling<br>Technology | CST 7076S |
|  | 1:5000 | Cell Signaling<br>Technology | CST 7074S |
| Mouse IgG | 1:5000 | Santa-Cruz | sc-3696 |

### RNA-sequencing

1 µg total RNA was used for NEBNext® Ultra II Directional RNA Library Prep Kit (New England Biolabs, Ipswich, MA, USA). Poly-A selection and cDNA synthesis were performed according to NEB protocol. The adaptors were diluted with a 1:30 ratio instead of the recommended 1:10 ratio. Size selection was performed using SPRI select beads. (Beckman Coulter, Indianapolis, IN, USA) with in-house calibration values. The cDNA was amplified with 22 cycles of PCR and 2 x150 paired-end sequencing was performed at a depth of 30 million reads at Novogene (Davis, Sacramento)

### RNA-seq data pre-processing

Human-HPV reference genome was constructed by combining HPV genome (https://pave.niaid.nih.gov/#search/search_database/locus_view/fetch?id=HPV16REF&format=Locus%20view&hasStructure=none) with human genome GRCh38.p13 from RefSeq(42). Quality control of paired-end raw sequencing data was conducted using FastQC-0.11.7 (43). Samples with adapter contamination were trimmed using Trimmomatic-0.38 (44). A second round of quality control was conducted by removing outlier samples based on clustering analysis of normalized gene expression matrix (also see the next section). Specifically, the distance measure for any pair of samples was defined as 1 minus the Pearson correlation of log-transformed gene expressions and was used for hierarchical clustering.

### Differential expression analysis

Raw sequencing data were aligned to human-HPV genome using Bowtie-1.2.2 (45). Gene-by-sample count matrix was calculated using RSEM-1.3.0(46). Genes with average expression less than 1 were filtered out. Median-by-ratio normalization (47) was conducted to get normalized expression matrix. Differential expression (DE) analysis was performed between JQ1+ and JQ1-samples using DESeq2-1.24.0 (48)under R version 3.6.0 (R Core Team, 2019). Genes with adjusted p-value less than or equal to 0.01 and absolute value of log2 fold change greater than or equal to 1 were selected as significant differentially expressed genes.

## HPV fusion analysis

Raw sequencing data were aligned to human-HPV genome using SpeedSeq-0.1.2 (49). Discordant reads are defined as read pairs not aligned to the reference genome with the expected distance or orientation. We focused on HPV-related discordant reads that one read mapped to HPV genome while the other mapped to human genome. Reads with mapping quality less than 10 were discarded. For the remaining HPV-related discordant reads, genome positions of those aligned to human genome were extracted as potential HPV fusion positions and mapped back to human genes to count the number of HPV fusion events for each gene. Gene annotation information was downloaded using R package biomaRt-2.40.4(50).

## Results

### Integrated HPV transcriptome displays sensitivity to pan-BET inhibitor, JQ1

BRD4, a member of the BET family of epigenetic co-regulators, has a known role in the HPV viral life cycle (33, 34, 51–57). Previous studies reported BRD4 to be overexpressed and correlated with poor prognosis in various solid cancers (58–61). To examine the expression profile of *BRD4* in HPV associated HNSCC, we evaluated the expression of BET genes involved in coregulating transcription using the publicly available curated TCGA dataset, UALCAN (62)which included 44 normal and 41 HPV associated HNSCC patients. As shown in **Figure 1A**, BRD1, BRD2-4 and BRD7-9 *w*ere significantly upregulated compared to normal samples in tumor samples classified to express p16, a surrogate for HPV detection and for presence for HPV DNA through *in situ* hybridization (p<0.05). In addition, as shown in **Figure 1B**, transcriptional co-regulators *BRD2, BRD3 and BRD4* (63, 64) were also significantly expressed (p<0.001) in tumors harboring HPV relative to normal tissue. Given these differences in expression, we hypothesized that HPV-associated tumors might exhibit a transcriptional reliance on BET proteins. This could enhance the transcription of oncogenic drivers, such as c-MYC, in head and neck tissues. Considering the differential expression of BRD4 between tumor and normal tissues, we proposed that targeted BET inhibitors might serve as a potential alternative to conventional genotoxic agents like cisplatin, which induces non-specific DNA cross-links (65, 66). To evaluate this hypothesis, we employed seven established integrated HPV-positive (+) HNSCC cell lines, differing in tumor origin and acquired epigenetic mutations. These cell lines differed in the location from which the original tumor was derived and in the epigenetic mutations that they acquired. To validate the presence of integrated HPV viral DNA, southern blot hybridization was performed to confirm viral integration in the host DNA. While in cell lines - 93VU147T, UPCI: SCC90 and UPC: SCC152 digested DNA released an 8kb viral DNA band, UD: SCC2, UM:SCC47, UM:SC104 and UPCI:SCC154 released higher molecular weight bands suggesting that viral DNA had either duplicated or undergone rearrangements upon integration into their respective host genomes. (Figure. S1).

**Figure 1:**
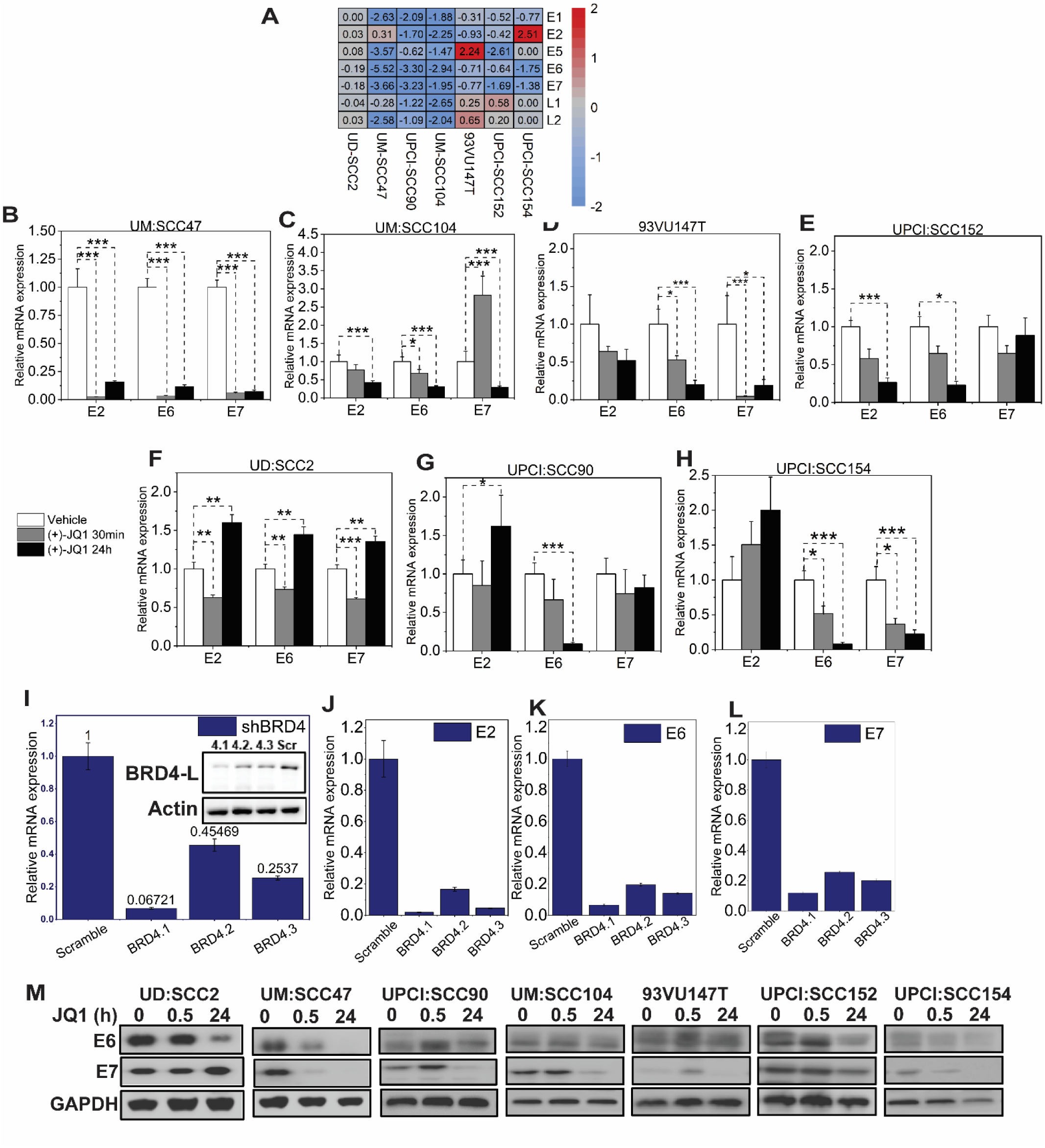
pan-BET inhibitor JQ1 treatment inhibits the growth of integrated HPV-associated head and neck squamous carcinoma cells (HNSCC) **(A)** TCGA data derived from UALCAN denotes BRD mRNA expression containing both bromodomains BD1 and BD2 associated with HPV status profiled through p16 and viral in situ hybridization from 41 patients was significantly higher compared to 44 normal patients. **(B)** Total read counts of all BRD RNAs retrieved from the TCGA database revealed higher expression in 80 patients compared to 40 normal patients. Y-axis represents transcripts per million (TPM), 25^th^ and 75^th^ percentile data, and Student’s t-test ** P<0.01, *** P<0.001 **(C)** IC_50_ values for seven HPV integrated HNSCC cell lines. Viable cells were determined using presto blue after treating cells with increasing concentration of JQ1 for 72 h. 72 h. Error bars denote the SDs of independent experiments using six wells per dose. **(D)** The anatomical location of the tumors with corresponding epigenetic drivers mutated in their coding regions was identified to be coregulated with BRD4. IC_50_ values are represented as the average ± SD (n =6). **(E)** Caspase 3/7 activity in seven integrated HPV-associated HNSCC cell lines from biological triplicate wells treated with 500 nM JQ1 (=)- and quantified through luminescence readouts, normalized to control (vehicle enantiomer JQ1 (-)-treated cells.

We then proceeded to determine the efficacy of pan-BET inhibitor, JQ1(+)-in these HPV-integrated cell lines. The bromodomain inhibitor JQ1(+) is known to selectively bind to the conserved bromodomains BD1 and BD2 of BRD4 with nanomolar affinities, thereby blocking its interaction with histones and consequently modulating transcription through protein-protein interactions (67). To assess the anti-proliferative potential of JQ1(+) in HPV-positive integrated cell lines, we examined its effect relative to the biologically inactive enantiomer, JQ1(-). Notably, the evaluated cell lines displayed half maximal inhibitory concentrations (IC50) ranging from 0.053µM to 0.786µM after 72 hours of JQ1(+) exposure (**Figure. 1C**, D, Figure S2). To ascertain the specificity of JQ1(+) in inhibiting cell growth, UM: SCC47 and UD: SCC2 cell lines, with similar growth profiles, were employed. Experiments at a concentration of 500 nM revealed that neither DMSO alone nor the inactive enantiomer JQ1(-) impacted cellular proliferation as assessed by the trypan blue assay (Figure. S3). To elucidate the mode of JQ1-induced growth inhibition, we undertook time-course experiments, focusing on apoptosis through measurement of Caspase3/7 activity. While there wasn’t a direct correlation between IC50 values and Caspase3/7 activity, all tested cell lines demonstrated a marked increase in apoptosis relative to vehicle controls (**Figure. 1E**). This observation was consistent with previous studies that JQ1 treatments increased apoptotic activity in lung adenocarcinoma cell lines (39).

### JQ1 treatment promotes E6 down-regulation compared to all viral genes

Given the role of viral DNA integration into the host genome in altering viral gene transcription, combined with the potential recruitment of BET proteins to integrated viral gene promoters, especially in head and neck squamous cell environment, we aimed to provide a comprehensive understanding of BET-protein mediated transcriptional landscape in these cell lines. A global RNA sequencing approach was employed to assess the consequences of BET protein-mediated transcriptional dysregulation. Seven integrated HPV-associated HNSCC cell lines were treated with JQ1 and subsequently analyzed at 24h in comparison with their respective enantiomer-treated controls at 500 nM. Viral reads were aligned to the human-HPV genome through discordant read mapping (49). Remarkably, barring UD:SCC2, there was a pronounced downregulation of viral oncogenes E6 and E7. Changes ranged from log2FC values of -5.52 to -1.52 for E6 and -3.66 to -1.62 for E7 (**Figure. 2A**).

**Figure 2:**
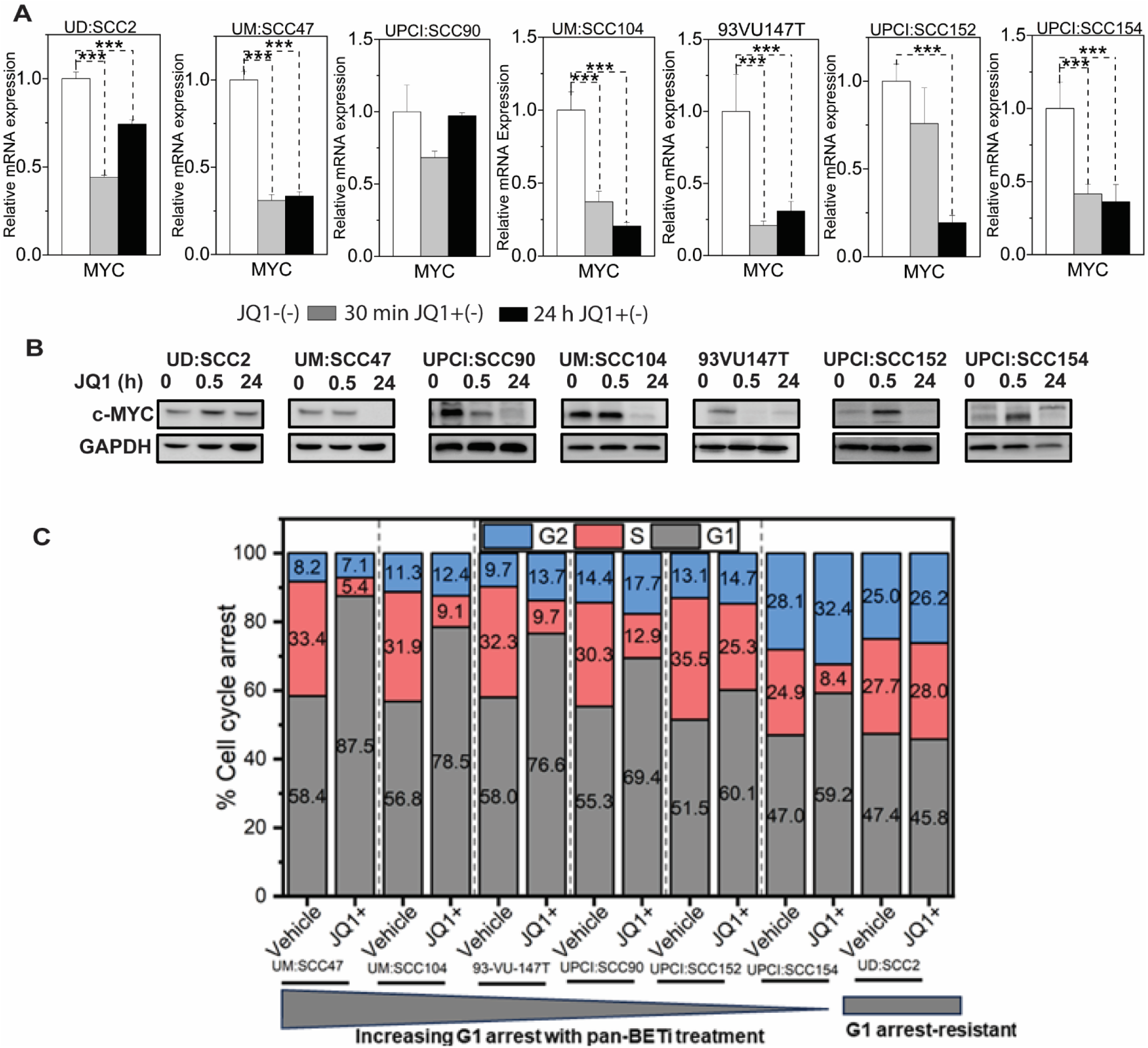
pan-BET inhibitor activity downregulates viral RNA and protein levels in selective HPV-associated HNSCC cells. **(A)** Discordant read mapping was performed to quantify viral read counts after RNA-sequencing experiments of seven integrated HPV-associated HNSCC cell lines treated with 500 nM JQ1 and its enantiomer control JQ1 (-)-at 24 hours. Data represented as heatmap and hierarchal clustering of log 2-fold changes in viral gene expression post-JQ1 treatment from DESeq2 analysis. The values inside each square represent log_2_ values with blue (down-regulated) and red (up-regulated). **(B-H)** Validation of RNA-sequencing viral gene expression was quantified through transcript levels of viral E2, E6, and E7 qRT-PCR in JQ1-treated cell lines at 500 nM concentration and normalized to vehicle JQ1 enantiomer at 30 minutes and 24 hours. RNA extraction was performed from three biological experiments, and gene expression was analyzed with three technical triplicates per experiment. Data are presented as the average ratio of viral E2, E6, and E7 RNA levels for each cell line relative to vehicle control (mean ± SEM). Asterisks denote the level of statistical significance - (*P < 0.05, **P < 0.01, ***P < 0.005; two-tailed t-test). **(I)** BRD4 knockdown phenocopies the effects of pan-BET inhibitor, JQ1, demonstrating downregulation of E2, E6, and E7 viral gene expression. Stable BRD4 knockdown was assessed in UM: SCC47 cell line by three independent shRNA clones, BRD4.1, 4.2, and 4.3, and quantified by qRT-PCR. Inset shows a western blot of Brd4 protein knockdown using anti-Brd4 antibodies. **(J-L)** Knockdown of BRD4 decreases E2, E6, and E7 expression in UM: SCC47 cell line. Data represented as the average ratio of viral gene expression relative to levels obtained from control scrambled shRNA UMSSCC47 cell line. shRNA decreases the expression. **(M)** Viral E6 and E7 protein levels in JQ1-treated sensitive integrated HPV-associated HNSCC cell lines. Cells treated with JQ1 and its enantiomer (vehicle) at 30 min and 24 h lysates were immunoblotted with anti-E6 and E7 antibodies; Gapdh serves as a loading control. Asterisks indicate cell lines that were probed using the same blotting membrane for viral E6 or E7 protein.

Previous reports have indicated a mechanism for E2-mediated repression that operates independently of Brd4, implying that distinct cellular components likely govern the transcriptional activation and suppression functions of E2 (68). Nonetheless, our data did not exhibit a consistent pattern of E2-dependent modulation of E6/E7 expression, even though the prevailing paradigm suggests a loss of E2 function upon integration. This observation was assessed across all the cell lines when quantifying the basal levels of these transcripts (Figure. S4). As seen in Figure.2A, specifically, when treated with JQ1 – in UPCI: SCC154 – a notable upregulation of the E2 gene was observed (2.51 log2 FC). Interestingly, this did not correlate with concurrent activation or repression of E6/E7 transcripts. Further, in UM: SCC104, UPCI: SCC90, 93VU147T and UPCI: SCC154 cell lines, a downregulation of the E2 gene was evident, with values of -2.25, -1.70, -0.93 and -0.42 log2 FC, respectively. This reduction did correlate with E6/E7 transcript downregulation. Additionally, while the UD: SCC2, UM: SCC47 and UPCI: SCC154 cell lines manifested elevated E2 regulation, E6/E7 expression was downregulated in these cell lines. Taken together, viral gene expression was modulated with JQ1 treatment, suggesting that BRD4 might actively participate in regulating E6/E7 viral genes independent of E2 function.

To validate the sequencing outcomes and account for potential discrepancies in paired-end read counts, qRT-PCR was performed. Additionally, the immediate effects of JQ1 on viral gene transcription were assessed at 30 minutes and compared to the 24-hour time point using qRT-PCR, to determine whether the overall inhibitory effect had an immediate effect on viral transcription. We noted varied responses of E2 transcription across the cell lines, but a more consistent downregulation of the E6 oncoprotein. E7 downregulation was observed in specific cell lines, but not uniformly across the panel. Our observations revealed differential transcriptional responses of the E2 gene across the cell lines studied, while the downregulation of the E6 oncoprotein was more consistent. Specifically, E2 expression decreased in four of the seven cell lines: UM: SCC47, UM: SCC104, 93VU147T, and UPCI: SCC152 (**Figure 2B**, D, E, and F). In contrast, UD: SCC2, UPCI: SCC90, and UPCI: SCC154 showed elevated E2 expression levels, though these were not statistically significant (**Figure 2A**, C, G). It’s worth noting that UPCI: SCC154, despite displaying a 2-fold increase in E2 expression, presented low absolute E2 levels (with a Cq value of 35), hinting at minimal or even absent E2 expression. In our study, six out of the seven cell lines showed a significant decrease in the E6 viral oncogene expression (**Figure 2** B-E,G,H). Moreover, in comparison to basal levels of E7, UM:SCC47, UM:SCC104, 93VU147T and UPCI: SCC154 showed significant downregulation of the viral oncoprotein (**Figure. 2** B, C, D, H). However, the remaining 3 cell lines (UD:SCC2, UPCI: SCC90 and UPCI: SCC152; (**Figure. 2** E,F,G) did not demonstrate significant downregulation of E7 compared to its basal levels. Further, examination of the functional characteristics of E6 and E7 viral oncogene expression revealed reduced protein levels of E6 in all cell lines except UPCI: SCC90, UMM: SCC104 and 93VU147T and reduced protein levels of E7 in all cell lines except UD: SCC2 at 24 h (**Figure. 2** M). We could not verify E2 protein levels due to the unavailability of reliable antibodies. While our findings indicated that JQ1 effectively inhibited viral gene expression, it was crucial to identify the role of BRD4 in this regulatory mechanism. To validate the involvement of BRD4 in governing E2, E6, and E7 expression, we employed a stable shRNA-mediated BRD4 knockdown in UM:SCC47 cells. The resulting downregulation of these viral genes upon BRD4 suppression, as shown in **Figure 2 I-L**, underscores BRD4’s integral role in modulating their expression. This not only established that JQ1 chemical inhibition was phenocopied with BRD4 knockdown but also strengthens the premise of BRD4’s pivotal role in viral gene transcriptional regulation.

### JQ1 treatment promotes MYC downregulation and modulates the p53-p21 axis

While the observed IC_50_ inhibitory effects on cellular survival may primarily stem from JQ1-mediated suppression of the viral oncogenes E6 and E7, we considered the potential influence of the well-established interaction between Brd4 and the oncogene c-Myc transcription (50, 69). Prior studies in cervical cancer have elucidated that HPV16 E6 and E7 proteins augment c-Myc expression through direct interactions (70).

Furthermore, viral integration events have been shown to potentiate c-Myc expression via long-range interaction mechanisms (71). Therefore, we assessed the impact of JQ1 on c-Myc, a recognized target of BRD4, postulating it as an ancillary mechanism underlying JQ1’s anti-proliferative action. We posited that a reduction in c-Myc levels, induced by JQ1, could serve as a complementary approach in modulating anti-proliferative effects, especially in JQ1-sensitive cell lines with diverse viral integration sites as documented earlier (72).

Indeed, JQ1 has been shown to diminish c-Myc expression in both hematological malignancies and solid tumors, thereby exerting its antitumor properties (73). Notably, certain subsets of lung adenocarcinoma cells demonstrated JQ1-mediated proliferation inhibition, irrespective of the downregulation of c-Myc (39), thus providing a rationale for potentially selecting HPV(+) patients for personalized treatment.

In our study, a marked downregulation of c-MYC expression was evident in JQ1-responsive cell lines both at the 30-minute and 24-hour timepoints following JQ1 administration. Relative to the control group treated with the JQ1(-) enantiomer, there was a significant decrease in c-MYC mRNA expression in all examined cell lines, exhibiting a fold change range between 0.15 to 0.75, 24 hours post JQ1(+) treatment (**Figure. 3A**). Intriguingly, while UD:SCC2 cell line showed unaltered c-Myc protein levels after 24 hours of JQ1(+) exposure, the remaining six cell lines demonstrated reduced or undetectable c-Myc protein quantities (**Figure. 3B**). Given c-Myc’s well-established role in modulating cell cycle progression and apoptosis-related mechanisms (74), we subsequently assessed the impact of JQ1 on cell cycle dynamics. Through propidium iodide staining and subsequent flow cytometry analysis, we observed a pronounced G1 phase arrest at the 24-hour mark in all c-MYC downregulated cell lines: UM: SCC47 (87.5%), UM: SCC104 (78.5%), 93VU147T (76.6%), UPCI: SCC152 (60%), and UPCI: SCC154 (59.2%), when compared with the corresponding inactive enantiomer treatments. Strikingly, UD: SCC2 emerged as an outlier, as it revealed absence of G1 phase restriction with cell cycle distribution being G1 (45.7%), S (28%), and G2 (26.1%) phases, compared to its control (**Figure. 3C**). In summary, our data suggests a strong correlation between c-Myc downregulation and G1 phase cell cycle arrest, underscoring c-Myc’s potential as aa concurrent therapeutic target for BET inhibition in integrated HPV(+) HNSCCs.

**Figure 3:**
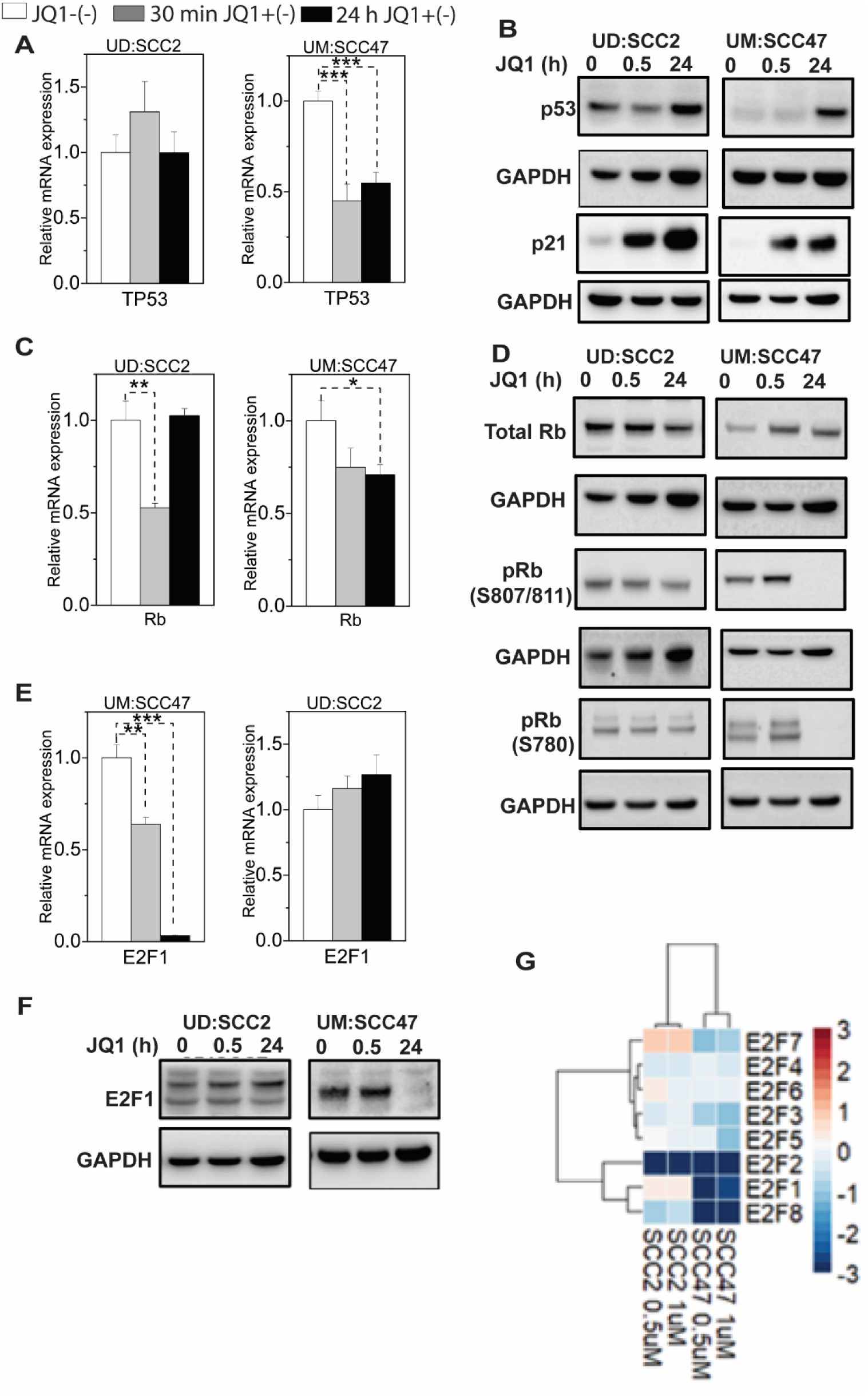
JQ1 treatment induces cell cycle arrest independent of c-Myc downregulation in six of the seven integrated HPV-associated HNSCC cell lines. **(A)** qRT-PCR evaluated c-MYC RNA levels in cells treated for 30 min and 24 hours with 500 nMJQ1 and its vehicle control. Mean fold change ±SEM (n=9) from three biological experiments. Asterisks denote the level of statistical significance - (*P < 0.05, **P < 0.01, ***P < 0.005; two-tailed t-test). **(B)** c-Myc protein levels in seven integrated HPV-associated HNSCC cell lines treated with 500 nM JQ1. Cells were treated with JQ1 and its inactive enantiomer for 30 min and 24 hours, lysed, and probed with anti-cMyc antibody. Gapdh serves as a loading control. **(C)** Cell-cycle analysis of seven integrated HPV-associated HNSCC cells treated with 500 nM for 24h and analyzed by propidium iodide flow cytometry. Stacked bar graphs show the percentage of cells in different cell cycle phases. The percentage of cells was calculated with Originlab software. Results represent mean values from three independent experiments.

### JQ1 treatment modulates p53-p21 axes

With underlying magnitude of G1 cell-cycle arrest mediated by JQ1 treatments in six of the seven cell lines, we wanted to determine if the inhibition of S-phase entry was mediated by additional targets controlled by the p53-BRD4-CDKN1A (p21) axis (75, 76). Recent studies have highlighted the potential of CDKN1A (p21) expression as a surrogate marker for HPV-related tonsillar squamous cell carcinoma, often indicating favorable patient outcomes. Specifically, a correlation was observed between CDKN1A (p21) and CDKN2A (p16) expression in HPV-associated laryngeal squamous cell carcinoma cases (77). Similarly, another study emphasized the significance of CDKN1A (p21), a tumor suppressor in the p53 pathway, as an indicator for HPV involvement in tonsillar squamous cell carcinoma (78, 79). Together, these findings suggest that CDKN1A (p21) can be a valuable biomarker not only for detecting HPV-associated cases but also for predicting patient prognosis.

In light of these reports, our study primarily centered on two contrasting cell lines: UM: SCC47, which exhibited the most pronounced G1 arrest, and UD:SCC2, which lacked this phenotype. Although RNA-seq read counts for TP53 did not reflect any pronounced JQ1-induced alterations in either cell line **(Figure. S5)**, RT-PCR results from UM: SCC47 showed a 50% decline in TP53 RNA levels post 24-hour JQ1 treatment (**Figure. 4A**). This decrease intriguingly corresponded with an increase in p53 protein concentrations (**Figure. 4B**). This increase in p53 could conceivably arise from the JQ1-mediated downregulation of E6 (**Figure. 2B**, **Figure. 2M**). However, despite the decrease of E6 protein in UD: SCC2, neither TP53 RNA nor p53 protein exhibited any substantial changes, post-JQ1 treatment, highlighting a plausible post-translational stability of p53 protein.

**Figure 4:**
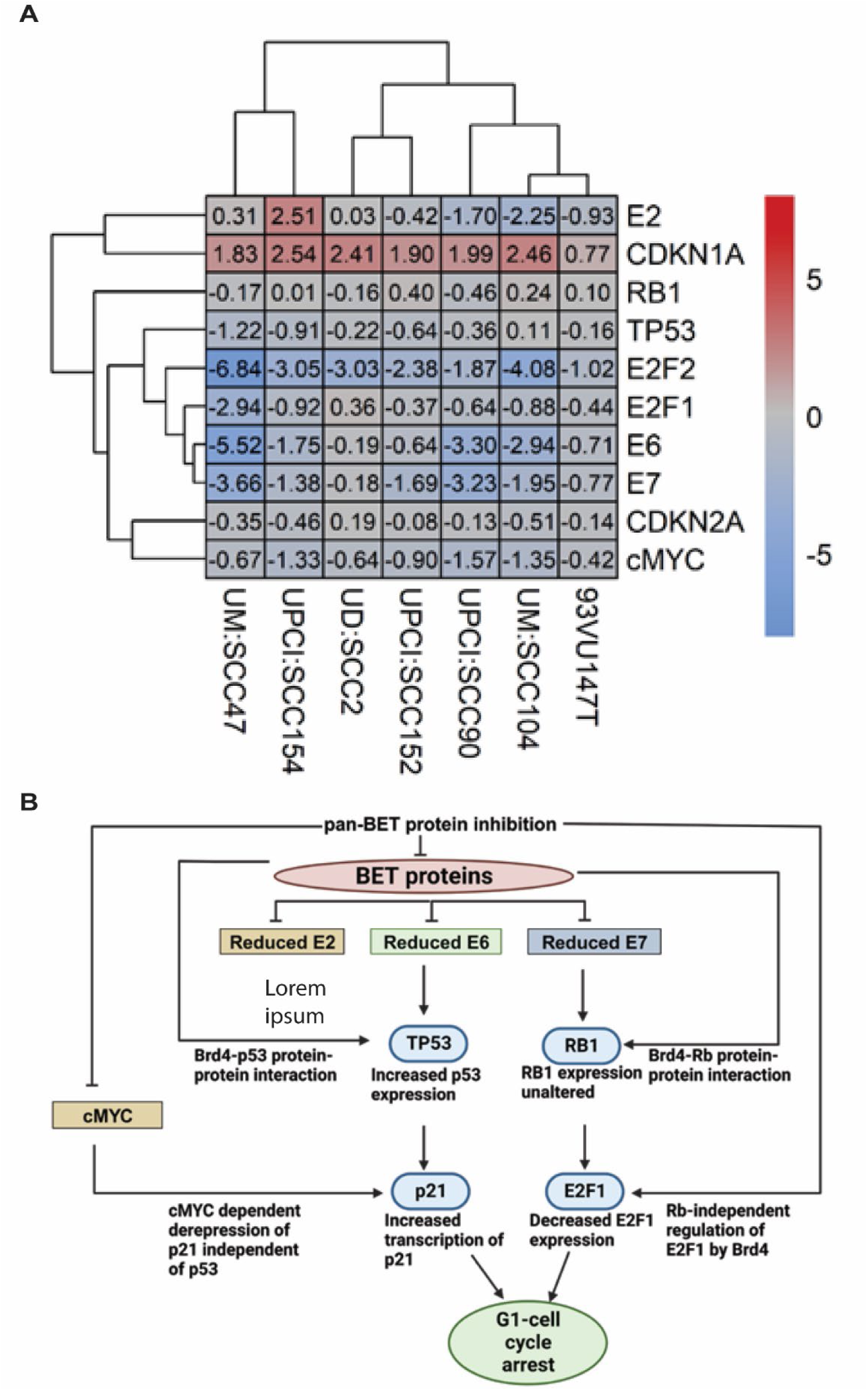
JQ1 treatments induce tumor suppressor p21 expression and down-regulate E2Fs expression. **(A)** qPCR analysis of TP53 expression in UMSCC47 (G1-arrest present) and UD: SCC2 (G1-arrest absent) cells treated with 500 nM JQ1 at 30 min and 24 h. **(B)** Western blot for total p53 and p21 in UD: SCC2 (G1-arrest absent) and UMSCC47 (G1-arrest present) cells treated for 30 min and 24 h with JQ1 and its inactive enantiomer. **(C)** qPCR analysis of RB1 expression in UD: SCC2 (G1-arrest absent) and UMSCC47 (G1-arrest present) cells under indicated time points with JQ1 at 500 nM. **(D)** Western blot of phosphorylated Rb relative to Total Rb in UD: SCC2 (G1-arrest absent) and UMSCC47 (G1-arrest present) cells under indicated treatments and time points. **(E)** qPCR analyses of the E2F1 gene in UD: SCC2 (G1-arrest absent) and UM: SCC47 (G1-arrest present) cells with JQ1 treatments at 30 min and 24 hours. **(F)** A heat map of an RNA-seq data of the log_2_ fold-change expression of the E2F family of genes that are either up-regulated (red) or down-regulated (blue) in UD: SCC2 and UM: SCC47 cells treated at 500 nM and 1µM JQ1 at 24 hours. For qPCR experiments, RNA from three biological experiments with triplicates (n=9) was used to perform statistical analyses, representing the average ratio of each target gene expression for each cell line relative to enantiomer control (mean ± SEM). RNA-seq experiments were conducted with three independent biological experiments.

### Relationship of c-Myc-p21 axis in response to JQ1 treatment

Upon further dissection into this complex network, we examined the downstream consequences of this p53 activation, particularly on the cyclin-dependent kinase (CDK) inhibitor p21(CDKN1A; WAF1/Cip1) -cMyc axis. Prior studies have demonstrated the ability of HPV-16 E7 to interact with Miz-1, subsequently inhibiting UV-induced p21^Cip1^ expression in C33A cervical (80). An alternative regulatory pathway involves the recruitment of c-Myc to the p21(WAF1/Cip1) promoter via Miz-1 (81), which was later shown that this interaction inhibits p53’s activation of p21 transcription (76). Our data indicate that the reduced/undetectable levels of both c-MYC and E6 proteins in UM:SCC47 likely facilitates the JQ1-induced upregulation of p21 (**Figure. 4B**). Conversely, UD:SCC2 cells treated with JQ1 presented a unique outlier: unaltered c-MYC and p53 protein levels and elevated p21 at 24 hours (**Figure. 4B**). This increase in p21, even in the presence of both c-MYC and p53, alludes to a possible dual regulation—both transcriptionally (via BET proteins) and through alternative mechanisms (82).

### E7-Rb-E2F Axis Dynamics Following JQ1 Treatment

With augmented p21 expression under the diverse cell-cycle arrest states observed in UD: SCC2 and UM:SCC47, we shifted our analyses towards understanding the network of the E7-Rb-E2F axis in relation to JQ1 treatment. Although prior literature elucidates the potential of E7-Rb interactions to degrade or inactivate pRb (83, 84), reports of endogenous E7-Rb inactivation or degradation within the context of an integrated viral HNSCC remain to be explored in detail. Moreover, the analysis of UD: SCC2 and UM: SCC47, particularly in terms of cell-cycle arrest, showed no marked changes in RB RNA expression levels **(Figure. S5),** further observed across all cell lines (**Figure. 2A**). This suggests that BRD4, despite its reported interaction with RB (85), might not be directly implicated in coregulating RB transcription.

Considering the established disruption of the Rb-E2F liaison by E7 (86), which consequentially stimulates E2F-mediated replication in S-phase (87), and the dependency of E2F-associated transcriptional programs on BET proteins (88, 89), we examined the impact of JQ1 treatment on the Rb-E2F axis in both UM: SCC47 and UD: SCC2 cells. While in UDSCC2, there was an initial downregulation of *RB* RNA to 50% at 30 min, the expression levels rebounded back at 24 h, comparable to untreated cells (**Figure. 4C**). Similarly, there was a marginal reduction of *RB* levels at 30 min and 24h by 25% in UM:SCC47 cells. Taken together, *RB* RNA levels were not significantly altered to infer whether JQ1 had a direct role in Rb downregulation. Previous reports in triple-negative breast cancer indicated that JQ1 treatments increased RB phosphorylation in a dose- and time-dependent manner with no alterations in E2F1 levels (90). This led us to address whether UD:SCC2 and UM:SCC47 cells would display different phosphorylation patterns post-JQ1 treatment. Interestingly, contrasting scenarios were observed: UD: SCC2 cells displayed unchanged total RB protein levels at both 30 min and 24 h points post-JQ1 treatment, in the presence of E7 protein (**Figure. 4D**), while concurrently displaying hyperphosphorylation at specific pRb (S807/811 and S780) sites. Conversely, UM: SCC47 cells displayed an increase in elevated total RB levels at the same time intervals, accompanied by reduced/undetectable levels of Rb phosphorylation (S807/811 and S780) at 24 hours (**Figure. 4D**). Crucially, the differential phosphorylation states observed in UD:SCC2 and UM: SCC47 cells, directly correlated to E2F1 protein release. The JQ1 treatment resulted in very significant downregulation to almost 95% of E2F1 RNA, and, correspondingly, the protein levels in UMSCC47 compared to UD: SCC2 **(Figure. 4E, F)**. These observations potentially hint at a possible transcriptional co-regulatory role of BET proteins in E2F1(91). Hence, we broadened our understanding on the effects of JQ1’s treatments on the broader E2F family members. Quantitative analysis of RNA expression through RNA-seq for the E2F members (E2F1-E2F8) in both UD: SCC2 and UM: SCC47 under two JQ1 concentrations (500 nM and 1µM) at 24 hours elucidated granular roles in JQ1-mediated G1 arrest. Remarkably, a significant downregulation of E2F1, 2, and 8 was observed in UM: SCC47 cells (**Figure. 4G**). Concurrently, other E2F genes exhibited similar downregulation compared to UD: SCC2 cells, with the most significant downregulation exclusively for E2F2 expression in UD: SCC2 and an increase in E2F7 (**Figure. 4G**). We quantified the RNA expression of all E2F members (E2F1-E2F8) in UD: SCC2 and UM: SCC47 at two different concentrations of JQ1-500 nM and 1µM for 24h to elucidate additional roles of JQ1-mediated G1 arrest. While E2F1, 2 and 8 were markedly downregulated in UMSCC47, other E2F gene members were also downregulated when compared to UDSCC2. Notably, the most significant downregulation was observed for E2F2 in UD: SCC2 with slight upregulation of E2F7 as well (**Fig 4G**). Although comprehensive regulatory aspects mediated by BET proteins encompassing E2F2-8 remain relatively obscure, our observations align with the notion of JQ1-driven downregulation of the E2F gene family, synergizing the G1 cell cycle arrest, and rendering a distinct physiological state in UM: SCC47 when contrasted with UD: SCC2 cells. Overall, in the context of HPV(+) HNSCC with integrated viral genes, our data highlight multiple concurrent therapeutic targets influenced by JQ1. In the context of HPV(+) HNSCC with randomly integrated viral sequences, our findings identify multiple concurrent targets that could have therapeutic implications when treated with JQ1. Collectively, these observations indicate unique response patterns to JQ1. Collectively, our results suggest distinct patterns of JQ1 response in part through downregulation of viral gene expression, which directly and indirectly impact the expression of cellular gene regulation in integrated HPV-HNSCC.

## Discussion

Our study reveals the intricate transcriptional responses induced by JQ1, a BET inhibitor, on key viral and cellular genes (Figure 5A). These findings have enabled us to construct a working model that sheds light on the regulatory role of BET proteins in the context of integrated HPV-associated HNSCC cell lines (**Figure 5B**). An important aspect of our study was to understand the well-documented interactions between Brd4 and the viral protein E2, with respect to its influence on E2 function (92–97) after viral integration While these studies have elegantly described the Brd4-E2 biology, it was unclear whether transcription of E2 gene would be disrupted in integrated viral HNSCC, due to the introduction of viral DNA breakpoints often associated with E2 gene (98, 99). Our analyses of the viral transcriptomes across all seven cell lines in this study suggested that E2 RNA expression remained abundant, as quantified through read counts by RNA sequencing **(Figure. S4A-B).** The only exception was UPCI: SCC154 cell line, where E2 expression was undetectable. These findings suggested that integration events did not result in the anticipated loss of the basal E2 open reading frame and JQ1 treatments modulated E2 expression. Since the impact of viral E2 gene disruption leads to loss of E2 function and can lead to increased expression of E6/E7 expression and contributes to carcinogenesis through uncontrolled cell proliferation, our quantification of the baseline expression of E2, E6/E7 expression did not indicate a clear ratio-metric pattern of E2 controlling E6 and E7 expression. However, it was notable, through shRNA BRD4 knockdown experiments, that BRD4 exerts a significant influence on the transcriptional regulation of E2, E6 and E7 expression (**Figure 5A**), and it phenocopies the effects of JQ1 inhibition. These findings parallel results observed in the 20861 cervical cell line, as reported by McBride et al for viral E6/E7 expression. This consistency in outcomes across different experimental systems underscores the potential therapeutic benefits of using BET inhibitors. While the mechanisms underlying the abundant RNA expression of episomal encoded viral genes could be likely influenced by factors such as the number of viral copies and the recruitment of cellular transcriptional machinery that includes BRD4, the expression of RNA in the integrated viral genomes is dependent on the genomic location of the integration event. Especially, in HPV-associated HNSCC genomes, the integrations are stochastic, potentially creating unique chromatin topologies consisting of super-enhancers that could modulate viral gene expression. We postulate that this could be the likely explanation for the heterogeneity of viral gene expression as identified through JQ1 treatments across the cell lines. Super-enhancers have been identified in the W12 cervical cell line 20861, where the viral genome had undergone tandem duplication of up to 30 copies at a single site, potentially regulating the amplitude of viral gene expression. Further investigations are underway in our laboratory to determine whether similar super-enhancers are formed when viral genomes integrate stochastically in HNSCC cells.

**Figure 5:**
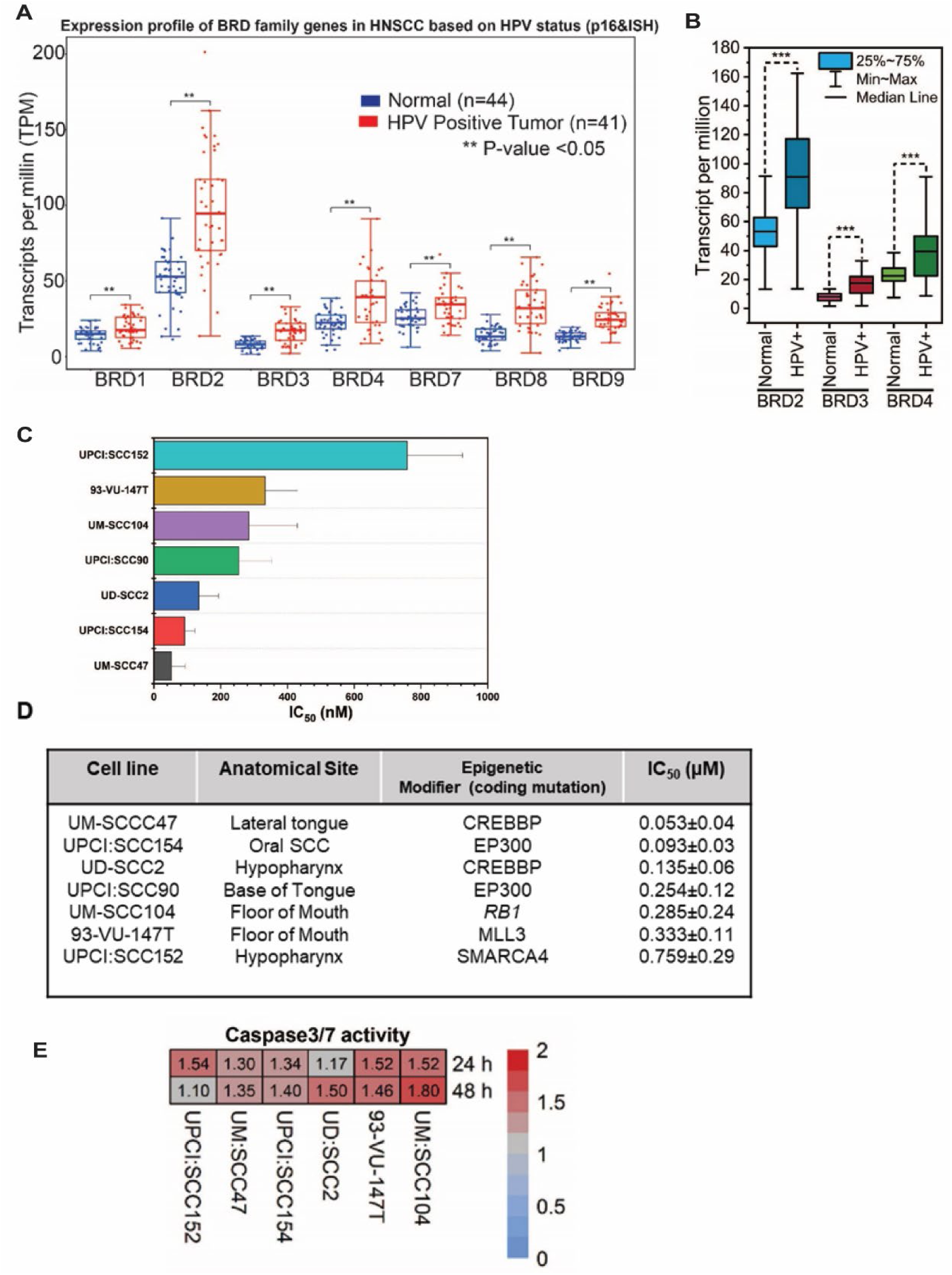
pan-BET inhibitor, JQ1 associated transcriptional landscape of key viral-cellular gene targets associated with cell-cycle progression in seven integrated HPV-associated HNSCC cells. **(A)** RNA-sequencing data from indicated seven cell lines treated with JQ1 and its enantiomer control at 24h. Discordant read-analyses for viral gene expression and cellular gene expression were performed from three independent biological replicates. A heat map represents a log_2_ fold change of viral and cellular genes with hierarchical clustering across seven cell lines. **(B)** Working model of pan-BET inhibitor, JQ1 treatment for integrated HPV-associated HNSCC cell lines suggesting an interplay of direct and indirect roles of BET proteins in regulating combined viral and cellular gene expression to induce a G1-arrest phenotype.

The consequence of viral gene expression in response to JQ1 resulted in G1-cell cycle arrest predominantly. Inhibiting BRD4 with JQ1 impacts cell-cycle progression (67, 100–102), indicating potential therapeutic roles in HPV-associated cancers. Of the seven tested cell lines, six showed G1-cell cycle arrest with JQ1. In contrast, the UD:SCC2 cell line did not undergo cell-cycle arrest. Instead, it exhibited hyperphosphorylation within 24 hours, in line with an absence of E7-Rb degradation, resulting in the continuous liberation of E2F1 for S-phase transcription. For the cell lines that did arrest in the G1 phase, JQ1 treatment led to reduced E2F expression, and Rb phosphorylation was absent by 24h. To summarize our G1 arrest findings post-JQ1 treatment: a) E2F gene family expression decreased, b) This decreased E2F expression triggered G1-arrest, and c) CDKs did not phosphorylate the freed E2Fs by 24h, an effect seen independently of JQ1’s influence. Our results align with earlier studies suggesting a possible interaction with or regulation between BRD4 and E2F genes, including super-enhancers (100, 101, 103, 104) in integrated HPV-associated HNSCC.

A further outcome of JQ1 treatments in these cell lines was the upregulation of p21 expression. Given that p53 is a significant inducer of p21, the interplay between viral E6 and p21 expression is particularly interesting. While G1-cell cycle arrest was observed in six of the seven tested cell lines with JQ1, we observed increased p21 expression across all seven cell lines. Notably, this observation has clinical implications since higher p21 expression in HPV-positive oropharyngeal cancers has identified a subset of these patients to have favorable prognosis (105) and p21 expression was absent or low in HPV-negative compared to HPV-positive cervical squamous cell carcinomas (106). From our baseline observations of p21 expression, these cell lines follow the classical notion for HPV-associated cancer is that given the known degradation of p53 by E6 protein, a direct consequence might be a decrease in p21 levels since p21 is a target of p53. However, our results suggest the rewiring of the viral–cellular signaling vis-à-vis E6-p53-p21 expression in response to JQ1 treatments. The increase in p21 expression is likely mediated by a decrease in E6 expression, which results in the absence or reduced degradation of p53. The result of this effect is an increase in p21 expression through p53 binding.

While reports of anti-leukemic activity was reported due to induction of p21 as one of the cellular responses to JQ1 treatment in acute myeloid leukemia (AML) cells(107), another study noted that JQ1 treatment led to growth arrest and apoptotic cell death and p21 expression was impacted, potentially implying its contribution to cell-cycle arrest, in triple-negative breast cancer(104). Taken together, it is worth noting that the response to JQ1 can be context-dependent, varying across cell or cancer types and particularly, our observations suggest that downregulating E6 is pivotal for modulating the p53-p21 expression. Given the established role of c-Myc in cellular proliferation and oncogenesis, its downregulation by JQ1 could have profound effects on cell-cycle progression.

Furthermore, if JQ1 concurrently downregulates the HPV E6 oncogene, it would significantly impact the cellular environment. The E6 protein has long been recognized for its ability to target and degrade the tumor suppressor p53, thus curtailing p53’s capacity to induce cell cycle arrest and apoptosis. Therefore, a reduction in E6 would lead to an increase in p53 levels. This stabilization of p53 would then prompt an upregulation of p21, a cyclin-dependent kinase inhibitor and a direct target of p53. Elevated levels of p21 can halt cell-cycle progression, thereby effectively putting a brake on cell proliferation.

Finally, we would like to postulate a regulatory framework that captures the relationships between c-Myc, E6, p53, and p21 expression in response to JQ1 treatment in the G1-arrested cells. If JQ1 concurrently downregulates E6 expression, it would significantly impact the viral-cellular environment. The E6 protein has long been recognized for its ability to target and degrade the tumor suppressor p53, thus curtailing p53’s capacity to induce cell cycle arrest and apoptosis. Therefore, a reduction in E6 would lead to an increase in p53 levels. This stabilization of p53 would then prompt an upregulation of p21, a cyclin-dependent kinase inhibitor and a direct target of p53. Elevated levels of p21 can halt cell-cycle progression, thereby effectively putting a brake on cell proliferation. In essence, the combined downregulation of c-Myc and E6 by JQ1 would create a synergistic cellular environment that is primed for cell-cycle arrest. With c-Myc levels reduced, the cell’s proliferative signals are diminished. Simultaneously, with E6 downregulated, p53 can accumulate and exert its tumor suppressor functions. This leads to an upsurge in p21, further enhancing the cell’s predisposition to arrest in the G1 phase of the cell cycle. In the case of UD:SCC2 cells that did not undergo G1-arrest, it is likely that either p53-mediated activation (108)or c-Myc-mediated repression (109, 110) can prevail and that both factors could be acting on the p21 promoter simultaneously, competing for transcriptional control. Taken together, it is noteworthy to observe that the overall effects of JQ1 treatments create various therapeutic intervention points both at the viral and cellular context.

In summary, our study presents a compelling argument for the potential therapeutic benefits of BET inhibitors in the context of integrated HPV-associated HNSCC. Further studies are ongoing to interrogate the role of BET proteins in super-enhancer formations in the context of random viral integrations, understanding if p21 expression is a double-edged sword for inducing cell-cycle arrest or creating a platform for senescence. Our findings have therapeutic implications for integrated HPV associated HNSCC cell lines which were sensitive to JQ1 treatment. With new generation of bromodomain (BD) inhibitors replacing JQ1, these drugs need to be explored as a viable targeted therapeutic strategy over standard of care nonspecific DNA targeting cisplatin.

## Author contributions

Conception and design, A.R., A.W., C.K., G.I.; data acquisition, A.R., D.S., A.W., K.N., D.L.; data analysis and interpretation (e.g., RNA-sequencing, computational analysis), Z.N., C.M., D.S., A.W., C.K., and G.I.; All other experiments, A.R., D.S., A.W.; manuscript writing. A.R. and G.I.

## Acknowledgements

The authors thank Drs. Kinjal Majumdar and Kavi Mehta for critical reading of the manuscript. This project was supported in part by NIH R21 CA267518-02 (G.I.), the University of Wisconsin Carbone Cancer Center Support Grant P30CA014520 and by the Specialized Program of Research Excellence (SPORE) program, through the National Cancer Institute (NCI), grant P50CA278595. The content is solely the responsibility of the authors and does not necessarily represent the official views of the NIH.

## Supplementary figures

### Nonlinear Curve Fit Dose Response Parameters

|  | Value | Standard Error | t-Value | Prob>\|t\| | Dependency |
| --- | --- | --- | --- | --- | --- |
| A1 | 42.49093 | 3.25572 | 13.05115 | 1.24718E5 | 0.48754 |
| A2 | 10398268 | 1399807 | 7.42836 | 3.06282E4 | 0.84385 |
| LOGx0 | -7.18672 | 0.18486 | -38.87645 | 1.93494E8 | 0.83835 |
| p | -1.66667 | 0.77999 | -2.13678 | 0.07649 | 0.75364 |
| span | 6149176 | 1533673 |  |  |  |
| EC20 | 1.49458E7 | 4.53401E8 |  |  |  |
| EC50 | 6.50552E8 | 2.76912E8 |  |  |  |
| EC80 | 2.83169E8 | 2.14424E8 |  |  |  |
| A1 | 47.62577 | 2.86894 | 16.60048 | 3.04803E6 | 0.43487 |
| A2 | 98.32513 | 5.57133 | 17.64842 | 2.12477E6 | 0.53333 |
| LOGx0 | -6.95525 | 0.12865 | -54.06292 | 2.68886E9 | 0.55615 |
| p | -1.66667 | 0.65585 | -2.54121 | 0.04401 | 0.53136 |
| span | 5069936 | 6.73198 |  |  |  |
| EC20 | 2.54674E7 | 8.61312E8 |  |  |  |
| EC50 | 1.10853E7 | 3.2838E8 |  |  |  |
| EC80 | 4.82516E8 | 2.53215E8 |  |  |  |
| A1 | 52.08379 | 2.1801 | 23.89059 | 3.53199E7 | 0.49136 |
| A2 | 94.55697 | 4.12771 | 22.90787 | 4.5335E7 | 0.61312 |
| LOGx0 | -6.79148 | 0.1189 | -57.11942 | 1.93422E9 | 0.5911 |
| p | -1.66667 | 0.54071 | -3.08235 | 0.0216 | 0.5907 |
| span | 4247318 | 5.06964 |  |  |  |
| EC20 | 3.71323E7 | 1.11008E7 |  |  |  |
| EC50 | 1.61628E7 | 4.42499E8 |  |  |  |
| EC80 | 7.03525E8 | 3.19445E8 |  |  |  |
| A1 | 59.52425 | 1.43547 | 41.46682 | 1.31561E8 | 0.28423 |
| A2 | 97.63684 | 2.81272 | 34.71266 | 3.80814E8 | 0.60298 |
| LOGx0 | -6.68143 | 0.08283 | -80.66333 | 2.44453E10 | 0.48042 |
| p | -1.66667 | 0.47598 | -3.50152 | 0.0128 | 0.4544 |
| span | 38.11259 | 3.35963 |  |  |  |
| EC20 | 4.78418E7 | 1.28869E7 |  |  |  |
| EC50 | 2.08244E7 | 3.97174E8 |  |  |  |
| EC80 | 9.06433E8 | 3.04773E8 |  |  |  |
| A1 | 16.85289 | 1.66196 | 10.14034 | 5.34922E5 | 0.14153 |
| A2 | 10186837 | 1580913 | 6.44364 | 6.61271E4 | 0.69777 |
| LOGx0 | -7.0341 | 0.16566 | -42.46106 | 1.14175E8 | 0.75703 |
| p | -1.66667 | 0.62101 | -2.6838 | 0.03635 | 0.57388 |
| span | 85.01549 | 16.09625 |  |  |  |
| EC20 | 2.12393E7 | 5.74289E8 |  |  |  |
| EC50 | 9.24495E8 | 3.52645E8 |  |  |  |
| EC80 | 4.0241E8 | 2.57667E8 |  |  |  |
| A1 | 68.48517 | 4.95197 | 13.82989 | 8.8954E6 | 0.40994 |
| A2 | 1094464 | 5.0427 | 21.70394 | 6.24656E7 | 0.57057 |
| LOGx0 | -6.51796 | 0.28336 | -23.00228 | 4.42407E7 | 0.33373 |
| p | -1.66667 | 1.578 | -1.05619 | 0.33154 | 0.59263 |
| span | 40.96123 | 8.06861 |  |  |  |
| EC20 | 6.97064E7 | 6.47238E7 |  |  |  |
| EC50 | 3.03415E7 | 1.97967E7 |  |  |  |
| EC80 | 1.32069E7 | 1.46452E7 |  |  |  |
| A1 | 74.91165 | 8.45115 | 8.86407 | 1.14692E4 | 0.80406 |
| A2 | 10750372 | 2.15514 | 49.88245 | 4.35383E9 | 0.35344 |
| LOGx0 | -5.82047 | 0.22501 | -25.86765 | 2.20081E7 | 0.88767 |
| p | -1.66667 | 1.24713 | -1.3364 | 0.22986 | 0.77963 |
| span | 32.59208 | 9.01 |  |  |  |
| EC20 | 3.4735E6 | 3.79727E6 |  |  |  |
| EC50 | 1.51193E6 | 7.83335E7 |  |  |  |
| EC80 | 6.58104E7 | 2.24638E7 |  |  |  |

**Statistics**
|  | UM-SCC-47 | UD-SCC-2 | UPCI-SCC-90 | 93VU147T | UPCI-SCC-154 | UM-SCC-104 | UPCI-SCC-152 |
| --- | --- | --- | --- | --- | --- | --- | --- |
| Number of Points | 10 | 10 | 10 | 10 | 10 | 10 | 10 |
| Degrees of Freedom | 6 | 6 | 6 | 6 | 6 | 6 | 6 |
| Reduced Chi-Sqr | 0.69464 | 0.32555 | 0.31803 | 0.46139 | 1.30944 | 0.72104 | 0.19722 |
| Residual Sum of Squares | 4.16786 | 1.9533 | 1.90819 | 2.76832 | 7.85662 | 4.32623 | 1.18331 |
| R-Square (COD) | 0.96798 | 0.98163 | 0.95816 | 0.95835 | 0.98063 | 0.88452 | 0.97049 |
| Adj. R-Square | 0.95197 | 0.97245 | 0.93723 | 0.93753 | 0.97094 | 0.82678 | 0.95574 |
| Fit Status | Succeeded(100) | Succeeded(100) | Succeeded(100) | Succeeded(100) | Succeeded(100) | Succeeded(100) | Succeeded(100) |

Fit Status Code : : Fit converged. Chi-Sqr tolerance value of 1E-9 was reached
|  |  | DF | Sum of Squares | Mean Square | F Value | Prob>F |
| --- | --- | --- | --- | --- | --- | --- |
| UM-SCC-47 | Regression | 4 | 3505.71854 | 876.42963 | 1261.69645 | 6.69166E-9 |
|  | Residual | 6 | 4.16786 | 0.69464 |  |  |
|  | Uncorrected Total | 10 | 3509.8864 |  |  |  |
|  | Corrected Total | 9 | 130.15589 |  |  |  |
| UD-SCC-2 | Regression | 4 | 1694.34777 | 423.58694 | 1301.14495 | 6.10211E-9 |
|  | Residual | 6 | 1.9533 | 0.32555 |  |  |
|  | Uncorrected Total | 10 | 1696.30107 |  |  |  |
|  | Corrected Total | 9 | 106.33643 |  |  |  |
| UPCI-SCC-90 | Regression | 4 | 1764.8204 | 441.2051 | 1387.29699 | 5.03577E-9 |
|  | Residual | 6 | 1.90819 | 0.31803 |  |  |
|  | Uncorrected Total | 10 | 1766.72859 |  |  |  |
|  | Corrected Total | 9 | 45.60214 |  |  |  |
| 93VU147T | Regression | 4 | 1525.05995 | 381.26499 | 826.34737 | 2.37625E-8 |
|  | Residual | 6 | 2.76832 | 0.46139 |  |  |
|  | Uncorrected Total | 10 | 1527.82826 |  |  |  |
|  | Corrected Total | 9 | 66.4735 |  |  |  |
| UPCI-SCC-154 | Regression | 4 | 1350.58848 | 337.64712 | 257.85689 | 7.70464E-7 |
|  | Residual | 6 | 7.85662 | 1.30944 |  |  |
|  | Uncorrected Total | 10 | 1358.4451 |  |  |  |
|  | Corrected Total | 9 | 405.51057 |  |  |  |
| UM-SCC-104 | Regression | 4 | 10685.41485 | 2671.35371 | 3704.87042 | 2.65067E-10 |
|  | Residual | 6 | 4.32623 | 0.72104 |  |  |
|  | Uncorrected Total | 10 | 10689.74108 |  |  |  |
|  | Corrected Total | 9 | 37.4633 |  |  |  |
| UPCI-SCC-152 | Regression | 4 | 3210.35867 | 802.58967 | 4069.54558 | 2.0003E-10 |
|  | Residual | 6 | 1.18331 | 0.19722 |  |  |
|  | Uncorrected Total | 10 | 3211.54198 |  |  |  |
|  | Corrected Total | 9 | 40.10021 |  |  |  |

At the 0.05 level, the fitting function is significantly better than the function $y=0$ .

**Figure.U S1.**
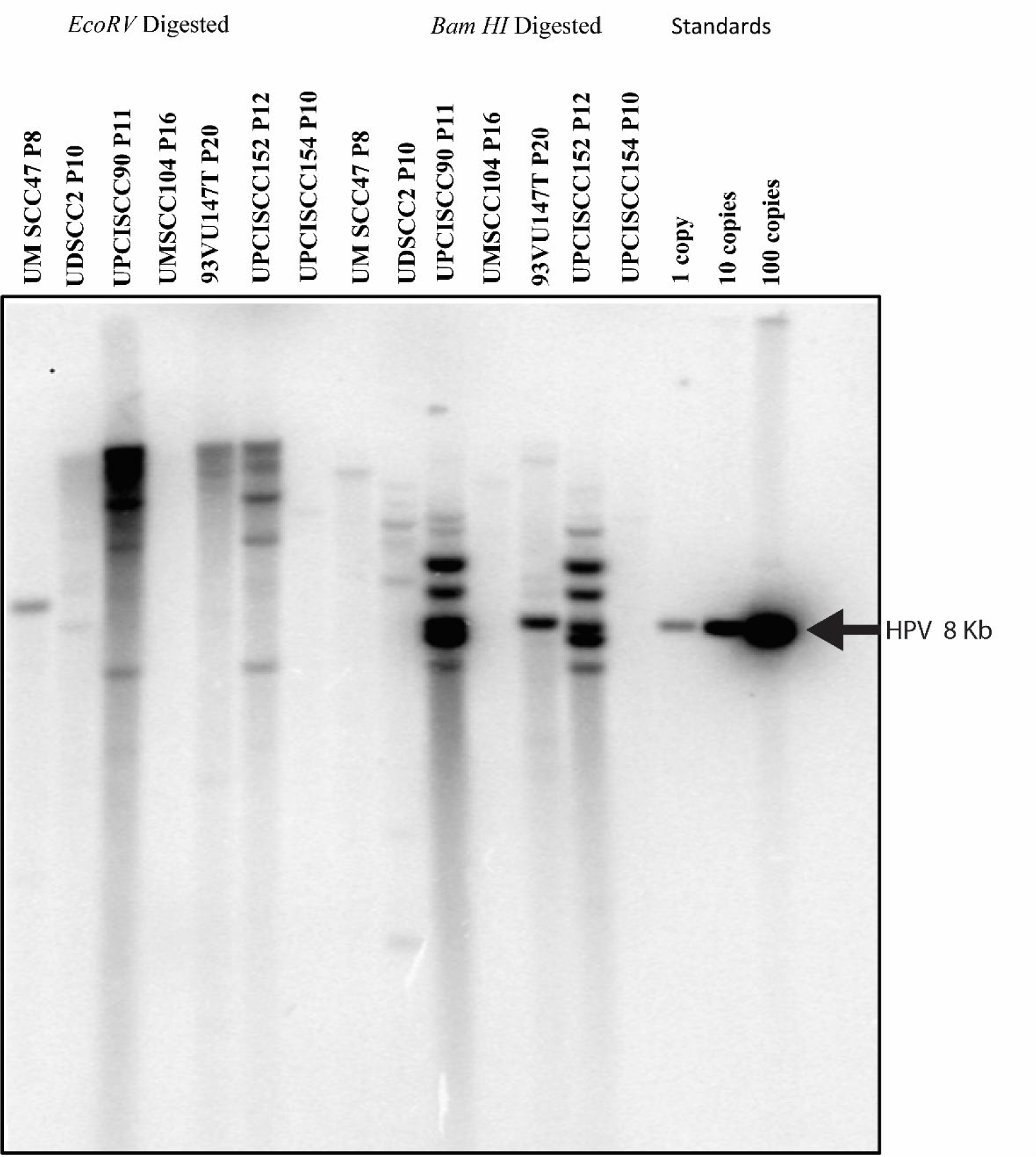
Southern blot hybridization of restriction digested of seven HPV cell lines. Control HPV ∼ 8 kb plasmid with copy number of 1, 10 and 100 was loaded to validate the presence of HPV DNA in these cancer cell lines. P indicates passage number.

**Figure. S2.**
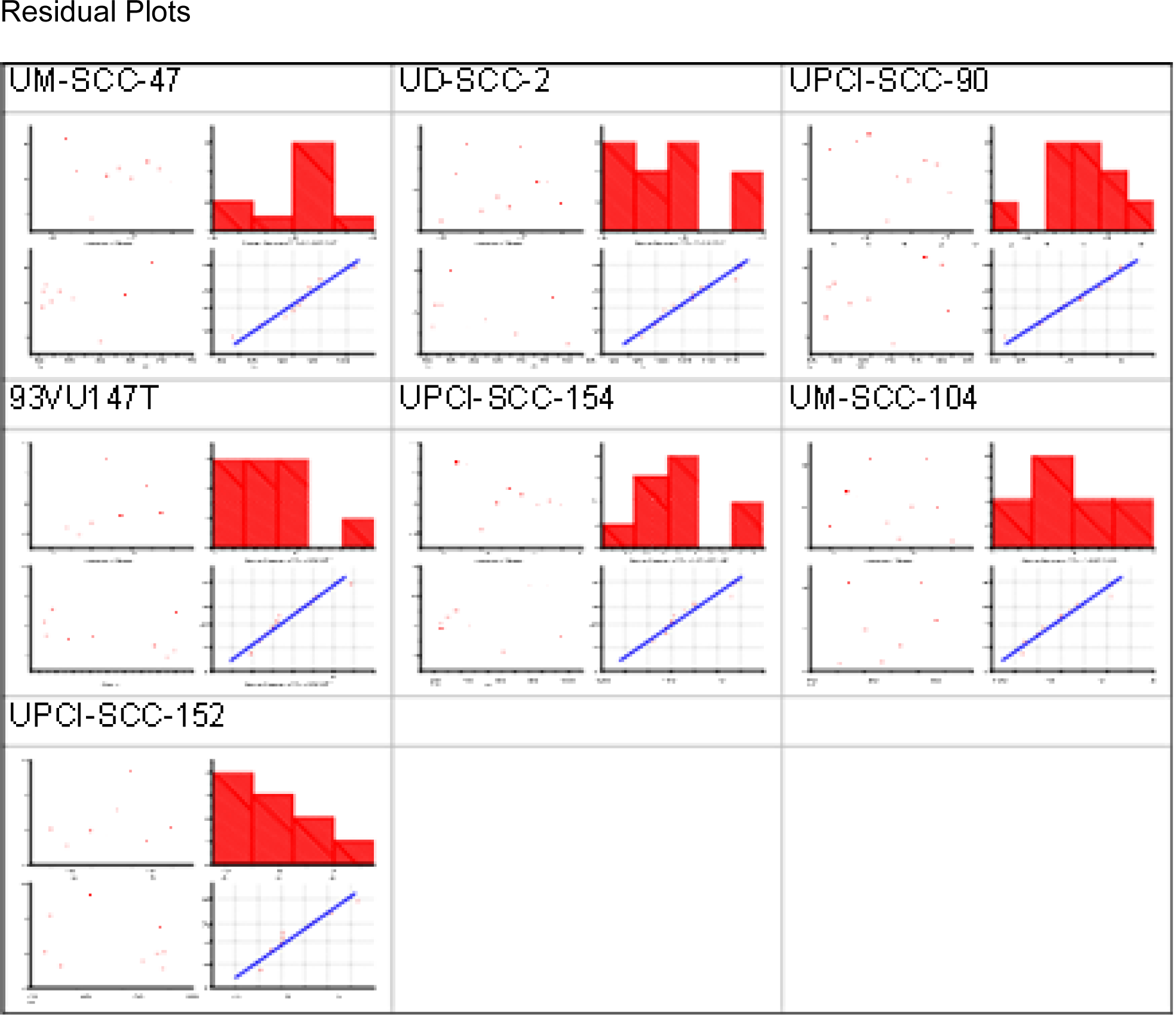
The IC50 values for JQ1 treatments for all cell lines were extracted from applying the Nonlinear Curve Fit using the Levenberg Marquardt algorithm using the DoseResp model. The residual fits confirm that the IC50 values were fitted to the equation y = A1 + (A2-A1)/(1 + 10^((LOGx0-x)*p))

**Figure. S3.**
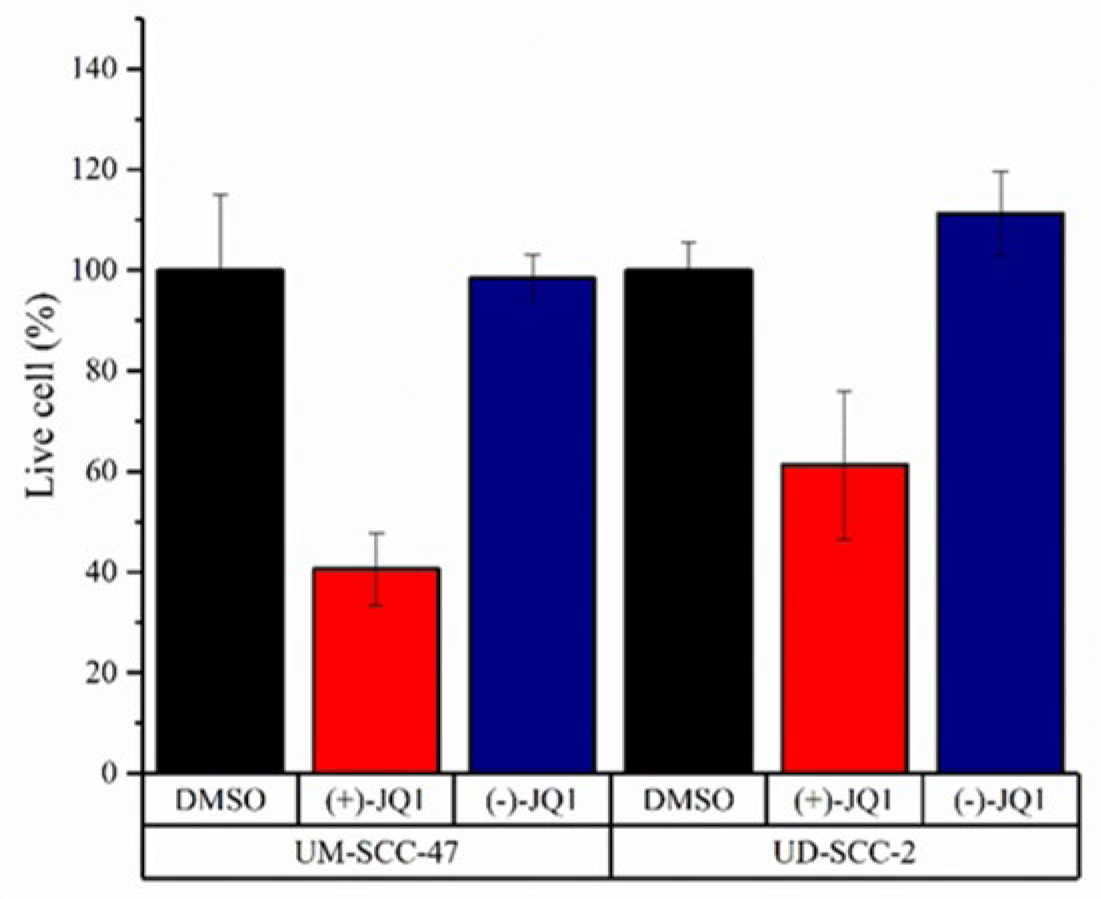
HPV-positive (UM-SCC-47, UD-SCC-2) head and neck cancer cell lines were treated with DMSO (vehicle), 0.5 μM (+)-JQ1 or (-)-JQ1 for 96 hours. The number of live cells in each condition was counted using a TC20 Biorad cell counter and trypan blue exclusion. The cell numbers were then normalized to the DMSO-treated group. With (+)-JQ1 treatment for 96 hours, the number of live cells was significantly reduced compared to vehicle control (DMSO). While the group treated with (-)-JQ1, a stereoisomer of (+)-JQ1, had a percentage of live cells similar to the vehicle control group. This suggests (-)-JQ1 has minimal effect on the proliferation of head and neck cancer cells at the concentration used in this study.

**Figure S4.**
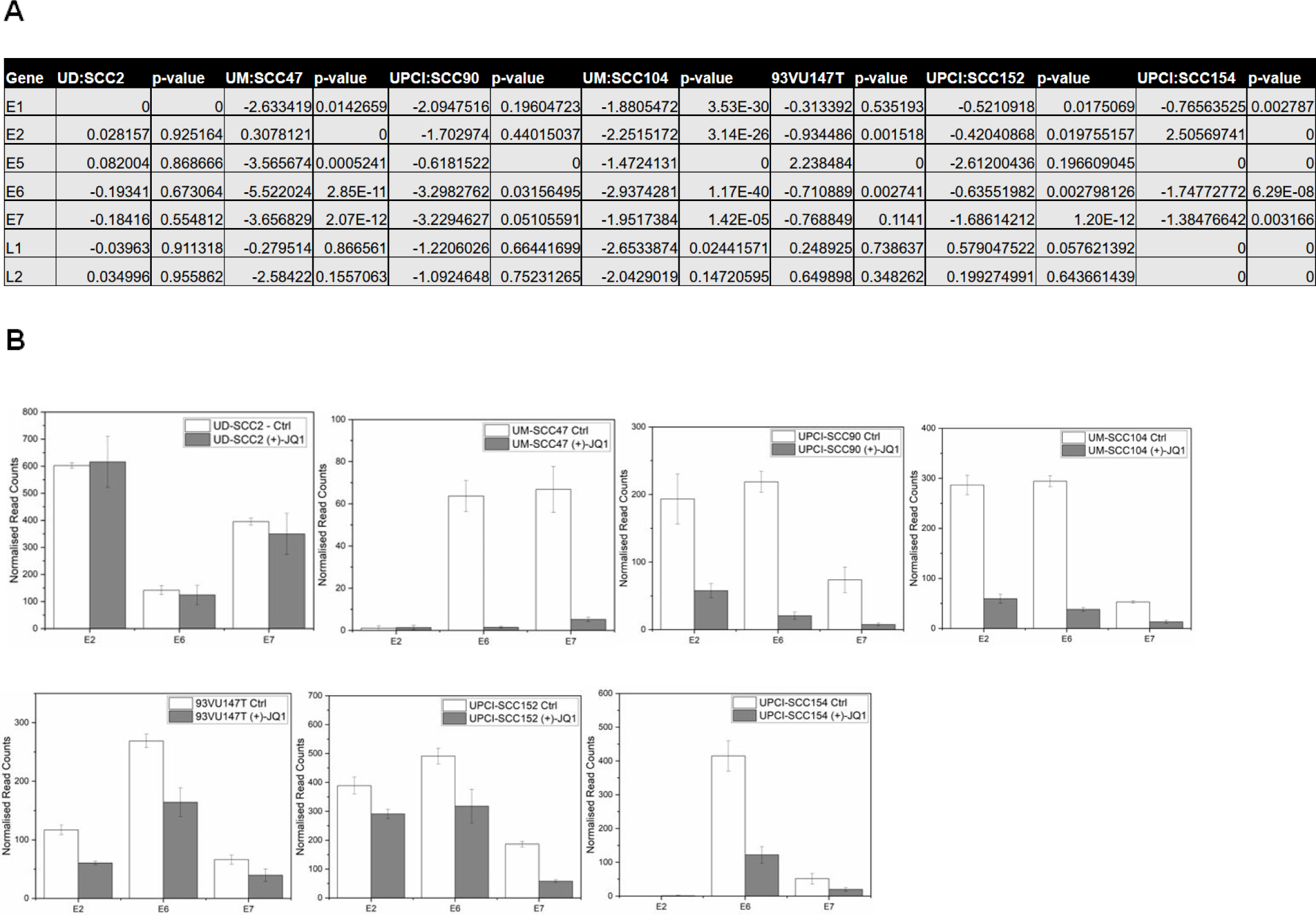
A. RNA-sequencing analyses of viral genes with log2fold change and its p-values. **B.** Normalized read counts of viral genes E2, E6, and E7 suggest the abundance of viral transcripts in seven HPV16-associated HNSCC cell lines.

**Figure S5.**
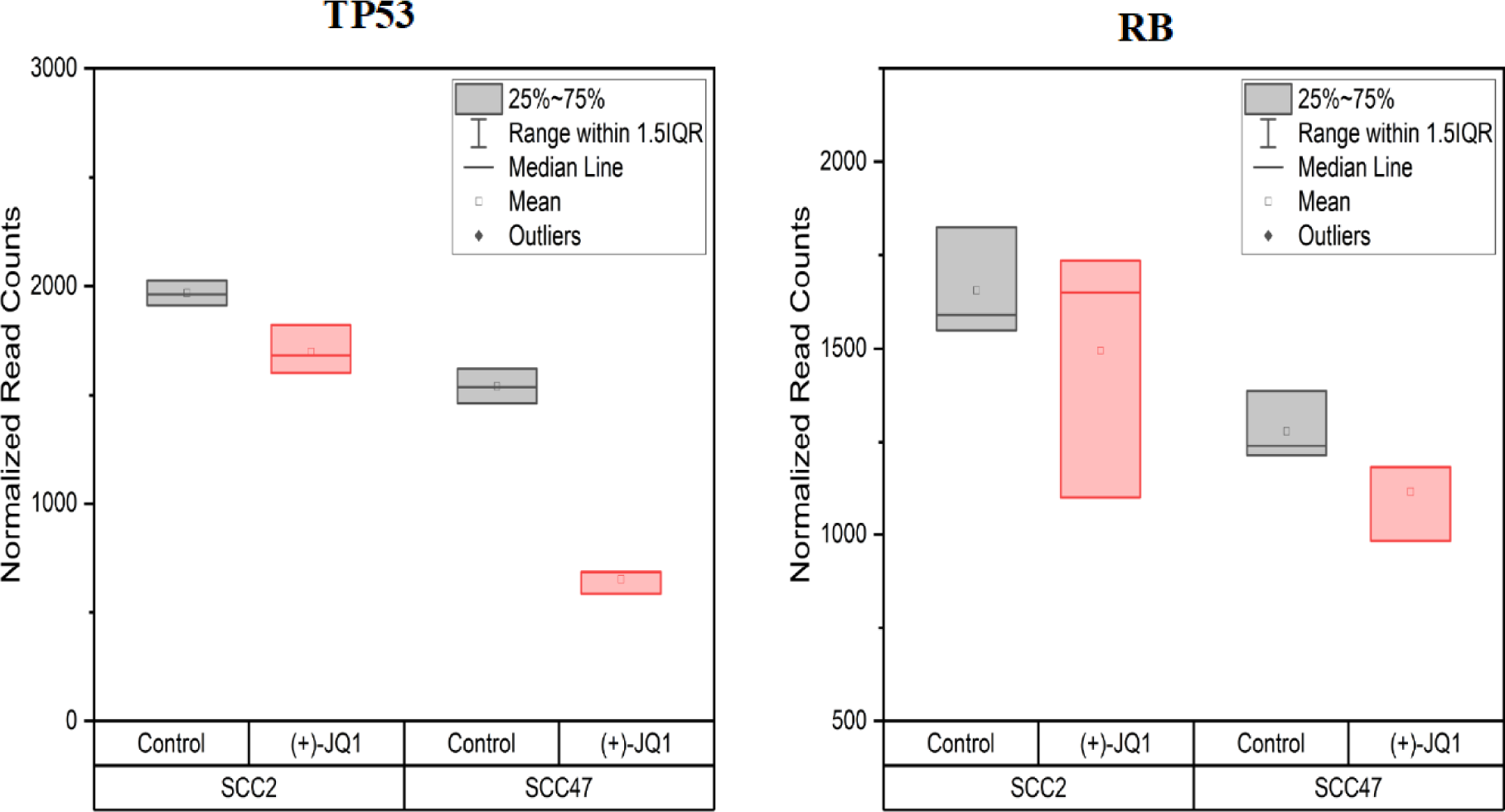
Box plot depicting normalized read counts of TP53 and RB from RNA-sequencing suggesting that despite viral E6 and E7 expression, both TP53 and RB RNA levels are abundantly expressed.

## References

1. Bravo IG, de Sanjosé S, Gottschling M. 2010. The clinical importance of understanding the evolution of papillomaviruses. Trends Microbiol 18:432–8.

2. Bernard HU, Burk RD, Chen Z, van Doorslaer K, zur Hausen H, de Villiers EM. 2010. Classification of papillomaviruses (PVs) based on 189 PV types and proposal of taxonomic amendments. Virology 401:70–9.

3. McBride AA. 2017. Mechanisms and strategies of papillomavirus replication. Biol Chem 398:919–927.

4. Longworth MS, Laimins LA. 2004. Pathogenesis of human papillomaviruses in differentiating epithelia. Microbiol Mol Biol Rev 68:362–72.

5. Pyeon D, Pearce SM, Lank SM, Ahlquist P, Lambert PF. 2009. Establishment of human papillomavirus infection requires cell cycle progression. PLoS Pathog 5:e1000318.

6. Bousarghin L, Touzé A, Sizaret PY, Coursaget P. 2003. Human papillomavirus types 16, 31, and 58 use different endocytosis pathways to enter cells. J Virol 77:3846–50.

7. Joyce JG, Tung JS, Przysiecki CT, Cook JC, Lehman ED, Sands JA, Jansen KU, Keller PM. 1999. The L1 major capsid protein of human papillomavirus type 11 recombinant virus-like particles interacts with heparin and cell-surface glycosaminoglycans on human keratinocytes. J Biol Chem 274:5810–22.

8. Burd EM. 2003. Human papillomavirus and cervical cancer. Clin Microbiol Rev 16:1–17.

9. zur Hausen H. 2009. Papillomaviruses in the causation of human cancers - a brief historical account. Virology 384:260–5.

10. Bergvall M, Melendy T, Archambault J. 2013. The E1 proteins. Virology 445:35–56.

11. McBride AA. 2013. The Papillomavirus E2 proteins. Virology 445:57–79.

12. Doorbar J. 2013. The E4 protein; structure, function and patterns of expression. Virology 445:80–98.

13. Regan JA, Laimins LA. 2008. Bap31 is a novel target of the human papillomavirus E5 protein. J Virol 82:10042–51.

14. Cortese MS, Ashrafi GH, Campo MS. 2010. All 4 di-leucine motifs in the first hydrophobic domain of the E5 oncoprotein of human papillomavirus type 16 are essential for surface MHC class I downregulation activity and E5 endomembrane localization. Int J Cancer 126:1675–82.

15. Hengstermann A, Linares LK, Ciechanover A, Whitaker NJ, Scheffner M. 2001. Complete switch from Mdm2 to human papillomavirus E6-mediated degradation of p53 in cervical cancer cells. Proc Natl Acad Sci U S A 98:1218–23.

16. Scheffner M, Werness BA, Huibregtse JM, Levine AJ, Howley PM. 1990. The E6 oncoprotein encoded by human papillomavirus types 16 and 18 promotes the degradation of p53. Cell 63:1129–36.

17. Helt AM, Galloway DA. 2003. Mechanisms by which DNA tumor virus oncoproteins target the Rb family of pocket proteins. Carcinogenesis 24:159–69.

18. Jones DL, Alani RM, Münger K. 1997. The human papillomavirus E7 oncoprotein can uncouple cellular differentiation and proliferation in human keratinocytes by abrogating p21Cip1-mediated inhibition of cdk2. Genes Dev 11:2101–11.

19. Wang JW, Roden RB. 2013. Virus-like particles for the prevention of human papillomavirus-associated malignancies. Expert Rev Vaccines 12:129–41.

20. Bedell MA, Hudson JB, Golub TR, Turyk ME, Hosken M, Wilbanks GD, Laimins LA. 1991. Amplification of human papillomavirus genomes in vitro is dependent on epithelial differentiation. J Virol 65:2254–60.

21. McBride AA. 2017. Oncogenic human papillomaviruses. Philosophical transactions of the Royal Society of London Series B, Biological sciences 372:20160273.

22. Dey A, Chitsaz F, Abbasi A, Misteli T, Ozato K. 2003. The double bromodomain protein Brd4 binds to acetylated chromatin during interphase and mitosis. Proceedings of the National Academy of Sciences 100:8758.

23. Yang Z, He N, Zhou Q. 2008. Brd4 recruits P-TEFb to chromosomes at late mitosis to promote G1 gene expression and cell cycle progression. Mol Cell Biol 28:967–76.

24. Dey A, Ellenberg J, Farina A, Coleman AE, Maruyama T, Sciortino S, Lippincott-Schwartz J, Ozato K. 2000. A bromodomain protein, MCAP, associates with mitotic chromosomes and affects G(2)-to-M transition. Mol Cell Biol 20:6537–49.

25. Mochizuki K, Nishiyama A, Jang MK, Dey A, Ghosh A, Tamura T, Natsume H, Yao H, Ozato K. 2008. The bromodomain protein Brd4 stimulates G1 gene transcription and promotes progression to S phase. J Biol Chem 283:9040–8.

26. Maruyama T, Farina A, Dey A, Cheong J, Bermudez VP, Tamura T, Sciortino S, Shuman J, Hurwitz J, Ozato K. 2002. A Mammalian bromodomain protein, brd4, interacts with replication factor C and inhibits progression to S phase. Mol Cell Biol 22:6509–20.

27. Drumond-Bock AL, Bieniasz M. 2021. The role of distinct BRD4 isoforms and their contribution to high-grade serous ovarian carcinoma pathogenesis. Mol Cancer 20:145.

28. Donati B, Lorenzini E, Ciarrocchi A. 2018. BRD4 and Cancer: going beyond transcriptional regulation. Molecular Cancer 17:164.

29. Li X, Baek G, Ramanand SG, Sharp A, Gao Y, Yuan W, Welti J, Rodrigues DN, Dolling D, Figueiredo I, Sumanasuriya S, Crespo M, Aslam A, Li R, Yin Y, Mukherjee B, Kanchwala M, Hughes AM, Halsey WS, Chiang CM, Xing C, Raj GV, Burma S, de Bono J, Mani RS. 2018. BRD4 Promotes DNA Repair and Mediates the Formation of TMPRSS2-ERG Gene Rearrangements in Prostate Cancer. Cell Rep 22:796–808.

30. Choi S, Bakkenist CJ. 2013. Brd4 shields chromatin from ATM kinase signaling storms. Sci Signal 6:pe30.

31. Floyd SR, Pacold ME, Huang Q, Clarke SM, Lam FC, Cannell IG, Bryson BD, Rameseder J, Lee MJ, Blake EJ, Fydrych A, Ho R, Greenberger BA, Chen GC, Maffa A, Del Rosario AM, Root DE, Carpenter AE, Hahn WC, Sabatini DM, Chen CC, White FM, Bradner JE, Yaffe MB. 2013. The bromodomain protein Brd4 insulates chromatin from DNA damage signalling. Nature 498:246–50.

32. Yan J, Diaz J, Jiao J, Wang R, You J. 2011. Perturbation of BRD4 protein function by BRD4-NUT protein abrogates cellular differentiation in NUT midline carcinoma. J Biol Chem 286:27663–75.

33. Dooley KE, Warburton A, McBride AA. 2016. Tandemly Integrated HPV16 Can Form a Brd4-Dependent Super-Enhancer-Like Element That Drives Transcription of Viral Oncogenes. mBio 7.

34. Warburton A, Redmond CJ, Dooley KE, Fu H, Gillison ML, Akagi K, Symer DE, Aladjem MI, McBride AA. 2018. HPV integration hijacks and multimerizes a cellular enhancer to generate a viral-cellular super-enhancer that drives high viral oncogene expression. PLoS Genet 14:e1007179.

35. Hnisz D, Abraham BJ, Lee TI, Lau A, Saint-Andre V, Sigova AA, Hoke HA, Young RA. 2013. Super-enhancers in the control of cell identity and disease. Cell 155:934–47.

36. Rataj O, Haedicke-Jarboui J, Stubenrauch F, Iftner T. 2019. Brd4 inhibition suppresses HPV16 E6 expression and enhances chemoresponse: A potential new target in cervical cancer therapy. Int J Cancer 144:2330–2338.

37. Nulton TJ, Kim NK, DiNardo LJ, Morgan IM, Windle B. 2018. Patients with integrated HPV16 in head and neck cancer show poor survival. Oral Oncol 80:52–55.

38. Balaji H, Demers I, Wuerdemann N, Schrijnder J, Kremer B, Klussmann JP, Huebbers CU, Speel EM. 2021. Causes and Consequences of HPV Integration in Head and Neck Squamous Cell Carcinomas: State of the Art. Cancers (Basel) 13.

39. Lockwood WW, Zejnullahu K, Bradner JE, Varmus H. 2012. Sensitivity of human lung adenocarcinoma cell lines to targeted inhibition of BET epigenetic signaling proteins. Proc Natl Acad Sci U S A 109:19408–13.

40. Iyer G, Wang AR, Brennan SR, Bourgeois S, Armstrong E, Shah P, Harari PM. 2017. Identification of stable housekeeping genes in response to ionizing radiation in cancer research. Sci Rep 7:43763.

41. Darzynkiewicz Z, Huang X, Zhao H. 2017. Analysis of Cellular DNA Content by Flow Cytometry. Current Protocols in Immunology 119:5.7.1-5.7.20.

42. O’Leary NA, Wright MW, Brister JR, Ciufo S, Haddad D, McVeigh R, Rajput B, Robbertse B, Smith-White B, Ako-Adjei D, Astashyn A, Badretdin A, Bao Y, Blinkova O, Brover V, Chetvernin V, Choi J, Cox E, Ermolaeva O, Farrell CM, Goldfarb T, Gupta T, Haft D, Hatcher E, Hlavina W, Joardar VS, Kodali VK, Li W, Maglott D, Masterson P, McGarvey KM, Murphy MR, O’Neill K, Pujar S, Rangwala SH, Rausch D, Riddick LD, Schoch C, Shkeda A, Storz SS, Sun H, Thibaud-Nissen F, Tolstoy I, Tully RE, Vatsan AR, Wallin C, Webb D, Wu W, Landrum MJ, Kimchi A, et al. 2016. Reference sequence (RefSeq) database at NCBI: current status, taxonomic expansion, and functional annotation. Nucleic Acids Res 44:D733–45.

43. Andrews S, Krueger, F., Seconds-Pichon, A., Biggins, F. & Wingett, S. 2015. FastQC: A quality control tool for high throughput sequence data. Babraham Institute.

44. Bolger AM, Lohse M, Usadel B. 2014. Trimmomatic: a flexible trimmer for Illumina sequence data. Bioinformatics 30:2114–2120.

45. Langmead B, Trapnell C, Pop M, Salzberg SL. 2009. Ultrafast and memory-efficient alignment of short DNA sequences to the human genome. Genome Biology 10:R25.

46. Li B, Dewey CN. 2011. RSEM: accurate transcript quantification from RNA-Seq data with or without a reference genome. BMC Bioinformatics 12:323.

47. Anders S, Huber W. 2010. Differential expression analysis for sequence count data. Genome Biology 11:R106.

48. Love MI, Huber W, Anders S. 2014. Moderated estimation of fold change and dispersion for RNA-seq data with DESeq2. Genome Biology 15:550.

49. Chiang C, Layer RM, Faust GG, Lindberg MR, Rose DB, Garrison EP, Marth GT, Quinlan AR, Hall IM. 2015. SpeedSeq: ultra-fast personal genome analysis and interpretation. Nat Methods 12:966–8.

50. Durinck S, Spellman PT, Birney E, Huber W. 2009. Mapping identifiers for the integration of genomic datasets with the R/Bioconductor package biomaRt. Nature Protocols 4:1184–1191.

51. Iftner T, Haedicke-Jarboui J, Wu SY, Chiang CM. 2017. Involvement of Brd4 in different steps of the papillomavirus life cycle. Virus Res 231:76–82.

52. Jang MK, Shen K, McBride AA. 2014. Papillomavirus genomes associate with BRD4 to replicate at fragile sites in the host genome. PLoS Pathog 10:e1004117.

53. McPhillips MG, Oliveira JG, Spindler JE, Mitra R, McBride AA. 2006. Brd4 is required for e2-mediated transcriptional activation but not genome partitioning of all papillomaviruses. Journal of virology 80:9530–9543.

54. Sakakibara N, Chen D, Jang MK, Kang DW, Luecke HF, Wu SY, Chiang CM, McBride AA. 2013. Brd4 is displaced from HPV replication factories as they expand and amplify viral DNA. PLoS Pathog 9:e1003777.

55. Lee AY, Chiang CM. 2009. Chromatin adaptor Brd4 modulates E2 transcription activity and protein stability. J Biol Chem 284:2778–2786.

56. Wu SY, Lee AY, Hou SY, Kemper JK, Erdjument-Bromage H, Tempst P, Chiang CM. 2006. Brd4 links chromatin targeting to HPV transcriptional silencing. Genes Dev 20:2383–96.

57. Gauson EJ, Wang X, Dornan ES, Herzyk P, Bristol M, Morgan IM. 2016. Failure to interact with Brd4 alters the ability of HPV16 E2 to regulate host genome expression and cellular movement. Virus Res 211:1–8.

58. Muhar M, Ebert A, Neumann T, Umkehrer C, Jude J, Wieshofer C, Rescheneder P, Lipp JJ, Herzog VA, Reichholf B, Cisneros DA, Hoffmann T, Schlapansky MF, Bhat P, von Haeseler A, Köcher T, Obenauf AC, Popow J, Ameres SL, Zuber J. 2018. SLAM-seq defines direct gene-regulatory functions of the BRD4-MYC axis. Science 360:800–805.

59. Andrieu G, Tran AH, Strissel KJ, Denis GV. 2016. BRD4 Regulates Breast Cancer Dissemination through Jagged1/Notch1 Signaling. Cancer Res 76:6555–6567.

60. Lu L, Chen Z, Lin X, Tian L, Su Q, An P, Li W, Wu Y, Du J, Shan H, Chiang C-M, Wang H. 2020. Inhibition of BRD4 suppresses the malignancy of breast cancer cells via regulation of Snail. Cell Death & Differentiation 27:255–268.

61. Asangani IA, Dommeti VL, Wang X, Malik R, Cieslik M, Yang R, Escara-Wilke J, Wilder-Romans K, Dhanireddy S, Engelke C, Iyer MK, Jing X, Wu YM, Cao X, Qin ZS, Wang S, Feng FY, Chinnaiyan AM. 2014. Therapeutic targeting of BET bromodomain proteins in castration-resistant prostate cancer. Nature 510:278–82.

62. Chandrashekar DS, Karthikeyan SK, Korla PK, Patel H, Shovon AR, Athar M, Netto GJ, Qin ZS, Kumar S, Manne U, Creighton CJ, Varambally S. 2022. UALCAN: An update to the integrated cancer data analysis platform. Neoplasia 25:18–27.

63. Jeanmougin F, Wurtz JM, Le Douarin B, Chambon P, Losson R. 1997. The bromodomain revisited. Trends Biochem Sci 22:151–3.

64. Winston F, Allis CD. 1999. The bromodomain: a chromatin-targeting module? Nat Struct Biol 6:601–4.

65. Corte-Rodríguez M, Espina M, Sierra LM, Blanco E, Ames T, Montes-Bayón M, Sanz-Medel A. 2015. Quantitative evaluation of cellular uptake, DNA incorporation and adduct formation in cisplatin sensitive and resistant cell lines: Comparison of different Pt-containing drugs. Biochem Pharmacol 98:69–77.

66. Wong E, Giandomenico CM. 1999. Current status of platinum-based antitumor drugs. Chem Rev 99:2451–66.

67. Filippakopoulos P, Qi J, Picaud S, Shen Y, Smith WB, Fedorov O, Morse EM, Keates T, Hickman TT, Felletar I, Philpott M, Munro S, McKeown MR, Wang Y, Christie AL, West N, Cameron MJ, Schwartz B, Heightman TD, La Thangue N, French CA, Wiest O, Kung AL, Knapp S, Bradner JE. 2010. Selective inhibition of BET bromodomains. Nature 468:1067–1073.

68. Schweiger MR, Ottinger M, You J, Howley PM. 2007. Brd4-independent transcriptional repression function of the papillomavirus e2 proteins. J Virol 81:9612–22.

69. Delmore Jake E, Issa Ghayas C, Lemieux Madeleine E, Rahl Peter B, Shi J, Jacobs Hannah M, Kastritis E, Gilpatrick T, Paranal Ronald M, Qi J, Chesi M, Schinzel Anna C, McKeown Michael R, Heffernan Timothy P, Vakoc Christopher R, Bergsagel PL, Ghobrial Irene M, Richardson Paul G, Young Richard A, Hahn William C, Anderson Kenneth C, Kung Andrew L, Bradner James E, Mitsiades Constantine S. 2011. BET Bromodomain Inhibition as a Therapeutic Strategy to Target c-Myc. Cell 146:904–917.

70. Peta E, Sinigaglia A, Masi G, Di Camillo B, Grassi A, Trevisan M, Messa L, Loregian A, Manfrin E, Brunelli M, Martignoni G, Palù G, Barzon L. 2018. HPV16 E6 and E7 upregulate the histone lysine demethylase KDM2B through the c-MYC/miR-146a-5p axys. Oncogene 37:1654–1668.

71. Shen C, Liu Y, Shi S, Zhang R, Zhang T, Xu Q, Zhu P, Chen X, Lu F. 2017. Long-distance interaction of the integrated HPV fragment with MYC gene and 8q24.22 region upregulating the allele-specific MYC expression in HeLa cells. Int J Cancer 141:540–548.

72. Akagi K, Li J, Broutian TR, Padilla-Nash H, Xiao W, Jiang B, Rocco JW, Teknos TN, Kumar B, Wangsa D, He D, Ried T, Symer DE, Gillison ML. 2014. Genome-wide analysis of HPV integration in human cancers reveals recurrent, focal genomic instability. Genome Research 24:185–199.

73. Shi J, Vakoc CR. 2014. The mechanisms behind the therapeutic activity of BET bromodomain inhibition. Mol Cell 54:728–36.

74. Amati B, Littlewood TD, Evan GI, Land H. 1993. The c-Myc protein induces cell cycle progression and apoptosis through dimerization with Max. The EMBO Journal 12:5083–5087.

75. Bretones G, Delgado MD, León J. 2015. Myc and cell cycle control. Biochimica et Biophysica Acta (BBA) - Gene Regulatory Mechanisms 1849:506–516.

76. Seoane J, Le HV, Massagué J. 2002. Myc suppression of the p21(Cip1) Cdk inhibitor influences the outcome of the p53 response to DNA damage. Nature 419:729–34.

77. Chernock RD, Wang X, Gao G, Lewis JS, Zhang Q, Thorstad WL, El-Mofty SK. 2013. Detection and Significance of Human Papillomavirus, CDKN2A(p16) and CDKN1A(p21) Expression in Squamous Cell Carcinoma of the Larynx. Modern Pathology doi:10.1038/modpathol.2012.159.

78. Kuo K-T, Hsiao C-H, Lin C-H, Kuo L-T, Huang S-H, Lin M-C. 2008. The Biomarkers of Human Papillomavirus Infection in Tonsillar Squamous Cell Carcinoma—molecular Basis and Predicting Favorable Outcome. Modern Pathology doi:10.1038/modpathol.3800979.

79. Hafkamp HC, Speel EJM, Haesevoets A, Bot FJ, Dinjens WNM, Ramaekers FCS, Hopman AHN, Manni JJ. 2003. A subset of head and neck squamous cell carcinomas exhibits integration of HPV 16/18 DNA and overexpression of p16INK4A and p53 in the absence of mutations in p53 exons 5–8. International Journal of Cancer 107:394–400.

80. Morandell D, Kaiser A, Herold S, Rostek U, Lechner S, Mitterberger MC, Jansen-Dürr P, Eilers M, Zwerschke W. 2012. The human papillomavirus type 16 E7 oncoprotein targets Myc-interacting zinc-finger protein-1. Virology 422:242–53.

81. Herold S, Wanzel M, Beuger V, Frohme C, Beul D, Hillukkala T, Syvaoja J, Saluz HP, Haenel F, Eilers M. 2002. Negative regulation of the mammalian UV response by Myc through association with Miz-1. Mol Cell 10:509–21.

82. Alani RM, Hasskarl J, Münger K. 1998. Alterations in cyclin-dependent kinase 2 function during differentiation of primary human keratinocytes. Mol Carcinog 23:226–33.

83. Boyer SN, Wazer DE, Band V. 1996. E7 protein of human papilloma virus-16 induces degradation of retinoblastoma protein through the ubiquitin-proteasome pathway. Cancer Res 56:4620–4.

84. Gonzalez SL, Stremlau M, He X, Basile JR, Münger K. 2001. Degradation of the retinoblastoma tumor suppressor by the human papillomavirus type 16 E7 oncoprotein is important for functional inactivation and is separable from proteasomal degradation of E7. J Virol 75:7583–91.

85. Ding D, Zheng R, Tian Y, Jimenez R, Hou X, Weroha SJ, Wang L, Shi L, Huang H. 2022. Retinoblastoma protein as an intrinsic BRD4 inhibitor modulates small molecule BET inhibitor sensitivity in cancer. Nature Communications 13:6311.

86. Chellappan S, Kraus VB, Kroger B, Munger K, Howley PM, Phelps WC, Nevins JR. 1992. Adenovirus E1A, simian virus 40 tumor antigen, and human papillomavirus E7 protein share the capacity to disrupt the interaction between transcription factor E2F and the retinoblastoma gene product. Proc Natl Acad Sci U S A 89:4549–53.

87. Cheng S, Schmidt-Grimminger DC, Murant T, Broker TR, Chow LT. 1995. Differentiation-dependent up-regulation of the human papillomavirus E7 gene reactivates cellular DNA replication in suprabasal differentiated keratinocytes. Genes Dev 9:2335–49.

88. Xu L, Chen Y, Mayakonda A, Koh L, Chong YK, Buckley DL, Sandanaraj E, Lim SW, Lin RY, Ke XY, Huang ML, Chen J, Sun W, Wang LZ, Goh BC, Dinh HQ, Kappei D, Winter GE, Ding LW, Ang BT, Berman BP, Bradner JE, Tang C, Koeffler HP. 2018. Targetable BET proteins- and E2F1-dependent transcriptional program maintains the malignancy of glioblastoma. Proc Natl Acad Sci U S A 115:E5086–e5095.

89. Peng J, Dong W, Chen L, Zou T, Qi Y, Liu Y. 2007. Brd2 is a TBP-associated protein and recruits TBP into E2F-1 transcriptional complex in response to serum stimulation. Mol Cell Biochem 294:45–54.

90. Shu S, Wu H-J, Ge JY, Zeid R, Harris IS, Jovanović B, Murphy K, Wang B, Qiu X, Endress JE, Reyes J, Lim K, Font-Tello A, Syamala S, Xiao T, Reddy Chilamakuri CS, Papachristou EK, D’Santos C, Anand J, Hinohara K, Li W, McDonald TO, Luoma A, Modiste RJ, Nguyen Q-D, Michel B, Cejas P, Kadoch C, Jaffe JD, Wucherpfennig KW, Qi J, Liu XS, Long H, Brown M, Carroll JS, Brugge JS, Bradner J, Michor F, Polyak K. 2020. Synthetic Lethal and Resistance Interactions with BET Bromodomain Inhibitors in Triple-Negative Breast Cancer. Molecular Cell 78:1096–1113.e8.

91. Xu L, Chen Y, Mayakonda A, Koh L, Chong YK, Buckley DL, Sandanaraj E, Lim SW, Lin RY-T, Ke X-Y, Huang M-L, Chen J, Sun W, Wang L-Z, Goh BC, Dinh HQ, Kappei D, Winter GE, Ding L-W, Ang BT, Berman BP, Bradner JE, Tang C, Koeffler HP. 2018. Targetable BET proteins- and E2F1-dependent transcriptional program maintains the malignancy of glioblastoma. Proceedings of the National Academy of Sciences 115:E5086.

92. McPhillips MG, Oliveira JGd, Spindler J, Mitra R, McBride AA. 2006. Brd4 Is Required for E2-Mediated Transcriptional Activation but Not Genome Partitioning of All Papillomaviruses. Journal of Virology doi:10.1128/jvi.01105-06.

93. Yigitliler A, Renner J, Simon C, Schneider M, Stubenrauch F, Iftner T. 2021. BRD4S Interacts With Viral E2 Protein to Limit Human Papillomavirus Late Transcription. Journal of Virology doi:10.1128/jvi.02032-20.

94. Zheng G, Schweiger M-R, Martínez-Noël G, Zheng L, Smith JA, Harper JW, Howley PM. 2009. Brd4 Regulation of Papillomavirus Protein E2 Stability. Journal of Virology doi:10.1128/jvi.00674-09.

95. Ilves I, Mäemets K, Silla T, Janikson K, Ustav M. 2006. Brd4 Is Involved in Multiple Processes of the Bovine Papillomavirus Type 1 Life Cycle. Journal of Virology doi:10.1128/jvi.80.7.3660-3665.2006.

96. Iftner T, Haedicke-Jarboui J, Wu SY, Chiang CM. 2017. Involvement of Brd4 in Different Steps of the Papillomavirus Life Cycle. Virus Research doi:10.1016/j.virusres.2016.12.006.

97. McBride AA, Jang M. 2013. Current Understanding of the Role of the Brd4 Protein in the Papillomavirus Lifecycle. Viruses doi:10.3390/v5061374.

98. Koskinen WJ, Chen RW, Leivo I, Mäkitie A, Bäck L, Kontio R, Suuronen R, Lindqvist C, Auvinen E, Molijn A, Quint W, Vaheri A, Aaltonen L-M. 2003. Prevalence and Physical Status of Human Papillomavirus in Squamous Cell Carcinomas of the Head and Neck. International Journal of Cancer doi:10.1002/ijc.11381.

99. Symer DE, Akagi K, Geiger HM, Song Y, Li G, Emde A-K, Xiao W, Jiang B, Corvelo A, Toussaint NC, Li J, Agrawal A, Ozer E, El-Naggar AK, Du Z, Shewale JB, Stache-Crain B, Zucker M, Robine N, Coombes KR, Gillison ML. 2022. Diverse tumorigenic consequences of human papillomavirus integration in primary oropharyngeal cancers. Genome Research 32:55–70.

100. Zuber J, Shi J, Wang E, Rappaport AR, Herrmann H, Sison EA, Magoon D, Qi J, Blatt K, Wunderlich M, Taylor MJ, Johns C, Chicas A, Mulloy JC, Kogan SC, Brown P, Valent P, Bradner JE, Lowe SW, Vakoc CR. 2011. RNAi screen identifies Brd4 as a therapeutic target in acute myeloid leukaemia. Nature 478:524–8.

101. Lovén J, Hoke Heather A, Lin Charles Y, Lau A, Orlando David A, Vakoc Christopher R, Bradner James E, Lee Tong I, Young Richard A. 2013. Selective Inhibition of Tumor Oncogenes by Disruption of Super-Enhancers. Cell 153:320–334.

102. Delmore JE, Issa GC, Lemieux ME, Rahl PB, Shi J, Jacobs HM, Kastritis E, Gilpatrick T, Paranal RM, Qi J, Chesi M, Schinzel AC, McKeown MR, Heffernan TP, Vakoc CR, Bergsagel PL, Ghobrial IM, Richardson PG, Young RA, Hahn WC, Anderson KC, Kung AL, Bradner JE, Mitsiades CS. 2011. BET bromodomain inhibition as a therapeutic strategy to target c-Myc. Cell 146:904–17.

103. Shi J, Wang Y, Zeng L, Wu Y, Deng J, Zhang Q, Lin Y, Li J, Kang T, Tao M, Rusinova E, Zhang G, Wang C, Zhu H, Yao J, Zeng Y-X, Evers BM, Zhou M-M, Zhou Binhua P. 2014. Disrupting the Interaction of BRD4 with Diacetylated Twist Suppresses Tumorigenesis in Basal-like Breast Cancer. Cancer Cell 25:210–225.

104. Shu S, Lin CY, He HH, Witwicki RM, Tabassum DP, Roberts JM, Janiszewska M, Jin Huh S, Liang Y, Ryan J, Doherty E, Mohammed H, Guo H, Stover DG, Ekram MB, Peluffo G, Brown J, D’Santos C, Krop IE, Dillon D, McKeown M, Ott C, Qi J, Ni M, Rao PK, Duarte M, Wu S-Y, Chiang C-M, Anders L, Young RA, Winer EP, Letai A, Barry WT, Carroll JS, Long HW, Brown M, Shirley Liu X, Meyer CA, Bradner JE, Polyak K. 2016. Response and resistance to BET bromodomain inhibitors in triple-negative breast cancer. Nature 529:413–417.

105. Weinberger PM, Yu Z, Haffty BG, Kowalski D, Harigopal M, Brandsma J, Sasaki C, Joe J, Camp RL, Rimm DL, Psyrri A. 2006. Molecular classification identifies a subset of human papillomavirus--associated oropharyngeal cancers with favorable prognosis. J Clin Oncol 24:736–47.

106. Milde-Langosch K, Riethdorf S, Kraus-Pöppinghaus A, Riethdorf L, Löning T. 2001. Expression of cyclin-dependent kinase inhibitors p16MTS1, p21WAF1, and p27KIP1 in HPV-positive and HPV-negative cervical adenocarcinomas. Virchows Arch 439:55–61.

107. Zhang H-T, Gui T, Sang Y, Yang J, Li Y-H, Liang G, Li T, He QY, Zha Z. 2017. The BET Bromodomain Inhibitor JQ1 Suppresses Chondrosarcoma Cell Growth via Regulation of YAP/p21/c-Myc Signaling. Journal of Cellular Biochemistry doi:10.1002/jcb.25863.

108. el-Deiry WS, Tokino T, Velculescu VE, Levy DB, Parsons R, Trent JM, Lin D, Mercer WE, Kinzler KW, Vogelstein B. 1993. WAF1, a potential mediator of p53 tumor suppression. Cell 75:817–25.

109. Nagashima M, Shiseki M, Pedeux RM, Okamura S, Kitahama-Shiseki M, Miura K, Yokota J, Harris CC. 2003. A novel PHD-finger motif protein, p47ING3, modulates p53-mediated transcription, cell cycle control, and apoptosis. Oncogene 22:343–50.

110. Prior IA, Harding A, Yan J, Sluimer J, Parton RG, Hancock JF. 2001. GTP-dependent segregation of H-ras from lipid rafts is required for biological activity. Nat Cell Biol 3:368–75.

